# Cohesin composition and dosage independently affect early development in zebrafish

**DOI:** 10.1101/2023.11.21.568176

**Authors:** Anastasia A. Labudina, Michael Meier, Gregory Gimenez, David Tatarakis, Sarada Ketharnathan, Bridget Mackie, Thomas F. Schilling, Jisha Antony, Julia A. Horsfield

## Abstract

Cohesin, a chromatin-associated protein complex with four core subunits (Smc1a, Smc3, Rad21 and either Stag1 or 2), has a central role in cell proliferation and gene expression in metazoans. Human developmental disorders termed “cohesinopathies” are characterised by germline mutations in cohesin or its regulators that do not entirely eliminate cohesin function. However, it is not clear if mutations in individual cohesin subunits have independent developmental consequences. Here we show that zebrafish *rad21* or *stag2b* mutants independently influence embryonic tailbud development. Both mutants have altered mesoderm induction, but only homozygous or heterozygous *rad21* mutation affects cell cycle gene expression. *stag2b* mutants have narrower notochords and reduced Wnt signaling in neuromesodermal progenitors as revealed by single cell RNA-sequencing. Stimulation of Wnt signaling rescues transcription and morphology in *stag2b*, but not *rad21* mutants. Our results suggest that mutations altering the quantity versus composition of cohesin have independent developmental consequences, with implications for the understanding and management of cohesinopathies.

**Summary Statement:** Viable zebrafish mutants show that cohesin complex quantity versus composition lead to different transcriptional and developmental outcomes in the early embryo.

## Introduction

Cohesin is a multiprotein ring-shaped complex that is highly conserved from yeast to humans. The vertebrate mitotic cohesin ring consists of two structural maintenance of chromosomes subunits, Smc1a, Smc3 and an *α*-kleisin subunit Rad21 (Huis et al., 2014; Nasmyth and Haering, 2009). In vertebrates Rad21 interacts with either one of two Stromalin subunits, Stag1 or Stag2, and collectively these subunits are necessary for cohesin’s association with DNA (Dorsett and Strom, 2012; Gruber et al., 2003; Horsfield et al., 2012). Additionally, Nipbl and Wapl modulate cohesin’s residency on chromatin: Nipbl facilitates loading of cohesin onto DNA (Ciosk et al., 2000), while Wapl facilitates its release (Kueng et al., 2006).

Cohesin is best known for its role in physically linking replicated sister chromatids to ensure the accurate transmission of genetic material to daughter cells during cell division (Nasmyth and Haering, 2009). In addition to mediating sister chromatin cohesion, the cohesin complex also functions to repair DNA double strand breaks (Hou et al., 2022; Sjögren and Nasmyth, 2001; Watrin and Peters, 2006). Loss of functional cohesin results in mitotic arrest and cell death (Cukrov et al., 2018; Horsfield et al., 2007; Percival et al., 2015). Only a small fraction of cohesin is necessary for sister chromatid cohesion (Gerlich et al., 2006) suggesting that the observed high levels of cohesin in certain non-dividing cell types has important non-cell cycle functions.

Cohesin also functions in three-dimensional (3D) genome organization and the regulation of gene expression (Ball et al., 2014; Cuadrado and Losada, 2020; Dorsett and Strom, 2012; Horsfield, 2023; Nishiyama, 2019; Zhu and Wang, 2019). Loop extrusion activity by cohesin organizes DNA into topologically associated domains (TADs) that constrain the regulation of gene expression (Davidson et al., 2019; Fudenberg et al., 2016; Hnisz et al., 2016; Krijger and de Laat, 2016; Sanborn et al., 2015). The CCCTC-binding factor, CTCF, acts as a barrier to cohesin and limits loop extrusion between convergent CTCF sites (Mayerova et al., 2020; Rao et al., 2014), leading to the overlap of cohesin and CTCF at TAD boundaries (Dixon et al., 2012; Parelho et al., 2008; Rao et al., 2014; Wendt et al., 2008). In addition, cohesin has gene regulatory functions that are independent of CTCF (Meier et al., 2018; Schmidt et al., 2010). Sites bound by cohesin but not CTCF are frequent at tissue-specific enhancers and promoters (Kagey et al., 2010). Intra-TAD loops formed by cohesin can regulate transcription by mediating enhancer-promoter contacts (Marsman and Horsfield, 2012; Ochi et al., 2020). However, only a subset of enhancer-promoter contacts and DNA looping events appear to depend on cohesin (Friman et al., 2023; Goel et al., 2023; Kane et al., 2022).

Germline cohesin insufficiency gives rise to a spectrum of multifactorial developmental disorders collectively known as ‘cohesinopathies’ (Ball et al., 2014; Horsfield et al., 2012). Typically, cohesinopathies result from heterozygous mutations in cohesin subunits or their regulators (Horsfield et al., 2012). Cohesinopathies are associated with developmental delay, a diverse range of developmental anomalies, and intellectual disability (Piché et al., 2019). The best known cohesinopathy is Cornelia de Lange syndrome (CdLS, MIM #122470), a multisystem disorder encompassing delayed growth, neurological and intellectual dysfunction, limb abnormalities and gastrointestinal defects (Ireland et al., 1993; Jackson et al., 1993; Liu et al., 2009; Opitz and Reynolds, 1985). Well over half of CdLS cases are caused by mutations in *NIPBL* (Krantz et al., 2004; Tonkin et al., 2004), with mutations in other cohesin-associated proteins accounting for a smaller subset of individuals with overlapping phenotypes. The specific presentation of CdLS varies according to the cohesin-associated protein affected by genetic changes (Cheung and Upton, 2015; Deardorff et al., 2020).

*RAD21* (MIM #606462) is among the five extensively studied genes associated with CdLS (Deardorff et al., 2012a; Deardorff et al., 2007; Krantz et al., 2004). Individuals with *RAD21* mutations display growth retardation, minor skeletal anomalies, and facial features that overlap with CdLS, but lack severe intellectual disabilities (Deardorff et al., 2012b). Mutations in *RAD21* are also linked with Mungan Syndrome (MIM #611376) (Mungan et al., 2003), sclerocornea (Zhang et al., 2019) and holoprosencephaly (Kruszka et al., 2019). Most *RAD21* mutations associated with cohesinopathy are truncations, missense mutations or in-frame deletions that are predicted to interrupt the interaction between RAD21 and SMC1A, SMC3, or STAG1/2 (Cheng et al., 2020; Krab et al., 2020). RAD21 physically bridges the SMC1A/SMC3 heads and facilitates the cohesin loading process, likely by controlling the amount that complexes with DNA (Sun et al., 2023). Therefore RAD21 abundance has potential to directly modulate the quantity of cohesin complexes on DNA and its mutation or deficiency would result in reduction in cohesin dose. Interestingly, the RAD21 protein must be intact for stable cohesin binding and looping at CTCF-CTCF sites, and must be present but not necessarily intact for looped contacts inside of CTCF domains (Liu and Dekker, 2022). Further supporting evidence suggests that the cohesion and loop extrusion activities of cohesin can be separated experimentally and that cohesin uses distinct mechanisms to perform these two functions (Nagasaka et al., 2023).

Individuals with STAG2 deficiency also display features of cohesinopathies (Cratsenberg et al., 2021; Kruszka et al., 2019; Mullegama et al., 2017; Mullegama et al., 2019; Soardi et al., 2017). Loss-of-function mutations in *STAG2* on the X chromosome are associated with Mullegama-Klein-Martinez syndrome (MKMS, MIM #301022) in females but only missense mutations are tolerated in males (Freyberger et al., 2021). Exome sequencing further established *STAG1* and *STAG2* variants in patients with cohesinopathy phenotypes as loss-of-function (Yuan et al., 2019) and recently, loss-of-function variants of *STAG2* have been categorized as X-linked cohesinopathies with features of CdLS (Mullegama et al., 2017; Soardi et al., 2017). For example an individual with a mosaic *STAG2* variant was described to have developmental delay, microcephaly, and hemihypotrophy of the right side (Schmidt et al., 2022). A distinctive cohesinopathy involving Xq25 microduplication that exclusively affects *STAG2* gives rise to moderate intellectual disability, speech delay and facial dysmorphism (Gokce-Samar et al., 2022). Additionally, some cases exhibit structural brain malformations consistent with holoprosencephaly (Cratsenberg et al., 2021; Kruszka et al., 2019; Mullegama et al., 2017; Soardi et al., 2017). Several molecular studies show that STAG1 and STAG2 paralogues have distinct roles in 3D genome organization, but overlapping roles in the cell cycle (Cheng et al., 2022; Cuadrado and Losada, 2020; Kojic et al., 2018; Porter et al., 2023; Remeseiro et al., 2012; Viny et al., 2019). Moreover, STAG subunits can be detected at specific locations on DNA independently of the rest of the cohesin complex (Pherson et al., 2019; Porter et al., 2023). Deficiency in STAG2 leads to the upregulation of STAG1 and the substitution of STAG1 for STAG2 in the cohesin complex such that *STAG2* mutation leads to altered cohesin composition (Adane et al., 2021; Bailey et al., 2021).

Dysregulated expression of multiple genes downstream of cohesin deficiency is thought to be the predominant cause of cohesinopathies (Dorsett and Krantz, 2009; Horsfield et al., 2007; Liu and Krantz, 2008; McNairn and Gerton, 2008; Muto et al., 2011). Because human cohesinopathies with different genetic causes present with diverse phenotypes, it is possible that cohesin subunits independently modulate the transcription function of cohesin during development. This idea has not yet been tested in the early embryo when the developmental changes in cohesinopathies are determined. In this study we compare the transcriptional and developmental consequences of depleting Rad21 with depletion of Stag2. Rad21 controls cohesin quantity on DNA (Sun et al., 2023) while Stag2 is thought to bind DNA independently and locate cohesin to enhancers (Kojic et al., 2018; Pherson et al., 2019; Porter et al., 2023). Therefore, we expect *stag2* mutants to interfere with cohesin’s gene expression functions without interfering with the cell cycle. Because *stag1b* and *stag2b* mutants are viable (Ketharnathan et al., 2020) and the effects of *rad21* deficiency are dose-dependent (Schuster et al., 2015), zebrafish offer a unique opportunity to investigate how cohesin complex quantity, versus cohesin complex composition, affects cell fate decisions in the early embryo (Muto and Schilling, 2017). To explore this question, we focused on the tailbud as a stem cell model.

The tailbud, located at the posterior end of the developing embryo, contains two populations of bipotent stem cells known as neuromesodermal progenitors (NMPs) and midline progenitor cells (MPCs) (Row et al., 2016; Steventon and Martin, 2022). These cells continuously divide and differentiate into neuroectoderm and mesoderm by activating cell type-specific transcription. By analysing *rad21* heterozygous and homozygous mutants (reflecting cohesin dose) and *stag2b* mutants (reflecting cohesin type), we compare how the amount and composition of the cohesin complex affect transcription in tailbud cells. We find that although *rad21* heterozygous mutants are viable and fertile, they exhibit altered expression of thousands of genes in the tailbud including cell cycle regulators, demonstrating that decreased cohesin dose affects both cell cycle and gene expression. In contrast, cell cycle gene expression is largely unaffected in *stag2b* homozygous mutants, which are also viable and fertile. However *stag2b* mutants show a unique narrowing of the midline mesodermal domain that forms the notochord.

Therefore, although both *rad21* and *stag2* cohesin mutants show deficiencies in mesoderm derived from NMPs and MPCs, the underlying molecular mechanisms are remarkably dissimilar. Rad21 deficiency blocks NMP differentiation leading to lack of mesodermal derivatives, while loss of *stag2* mutants causes NMPs to downregulate Wnt signaling leading to epithelial to mesenchyme (EMT) defects. Changes in phenotype and gene expression unique to *stag2* mutants are rescued by stimulation of Wnt signaling by GSK3 inhibition, to which Rad21-deficient embryos are impervious.

## Results

### Combined loss of zebrafish Stag1b and Stag2b phenocopies the null cohesin mutation *rad21*

Zebrafish have four Stag paralogues: Stag1a, Stag1b, Stag2a, and Stag2b. Individual *stag* mutant lines (except *stag2a*) were previously generated and are homozygous viable (Ketharnathan et al., 2020). To determine which paralogs are crucial for zebrafish development, we analysed the consequences of combining *stag1a* and *stag2b* as well as *stag1b* and *stag2b* mutants.

*stag1b^-/-^*; *stag2b^-/-^* double mutant embryos were developmentally delayed compared to wild type, and by ∼48 hours post-fertilization (hpf), mutant embryos had arrested in development presenting with small heads, pericardial oedema, upwards bending tails, and no blood circulation (Fig. 1B compared with A). This phenotype resembles *rad21^-/-^* mutant embryos, which die due to mitotic catastrophe (Fig. 1C) (Horsfield et al., 2007). In contrast, *stag1a^-/-^*; *stag2b^-/-^* embryos developed normally and are homozygous viable and fertile, although a small proportion (∼5%) of *stag1a^-/-^*; *stag2b^-/-^* embryos displayed hemorrhaging above the notochord at 48 hpf (Fig. S1A).

**Fig. 1.**
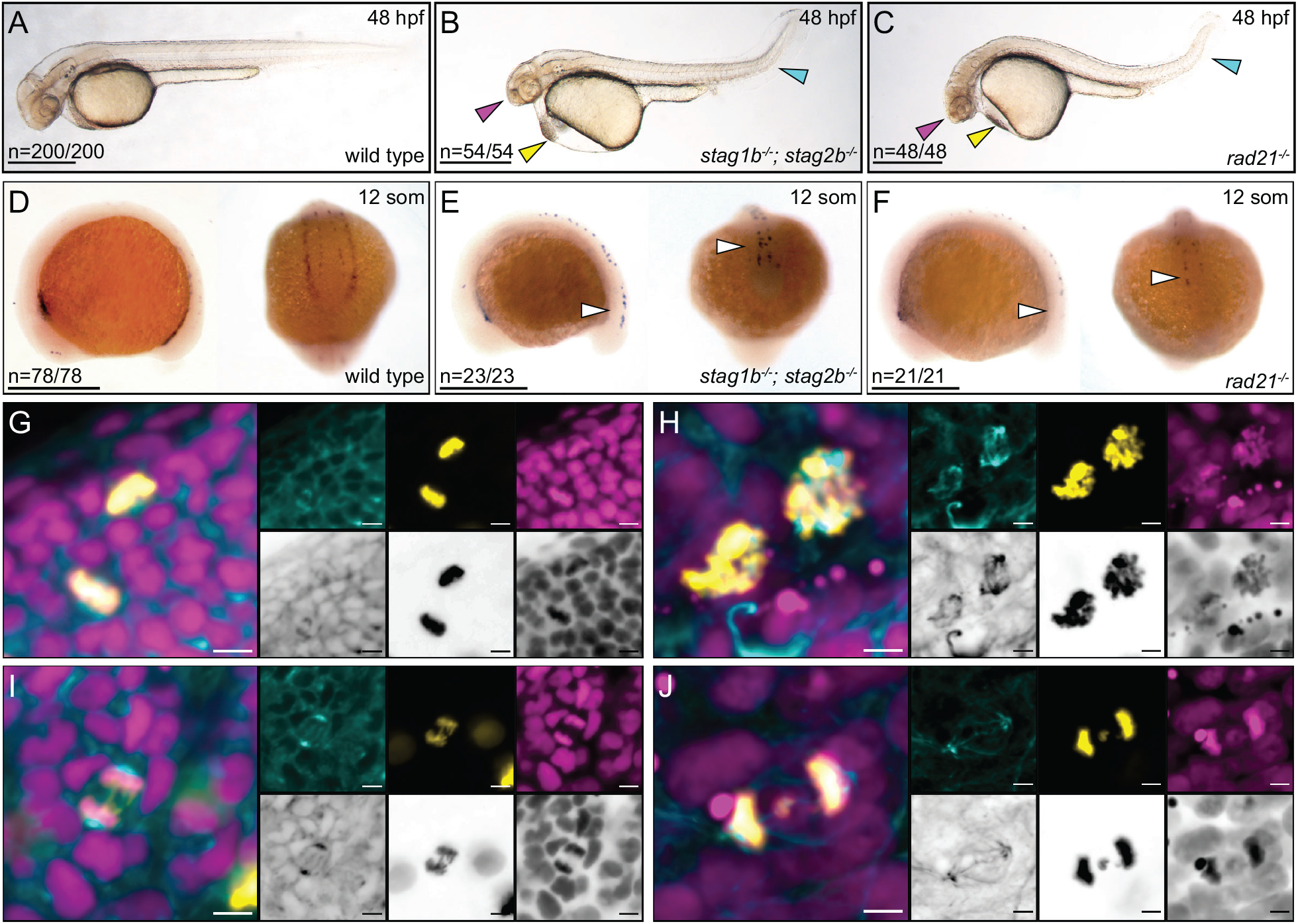
Combined loss of zebrafish Stag1b and Stag2b phenocopies the null cohesin mutation *rad21*. **(A-C)** Lateral views of representative wild type (A), *stag1b*^-/-^; *stag2b*^-/-^ (B) and *rad21*^-/-^ (C) embryos at 48 hours post-fertilization (hpf). Arrows indicate developmental anomalies: magenta for a small head, yellow for pericardial oedema, and cyan for a kinked tail. Scale bars are 500 μm. **(D-F)** Expression of *runx1* at 12 somites in wild type (D), *stag1b*^-/-^; *stag2b*^-/-^ (E) and *rad21*^-/-^ (F) embryos. Lateral and posterior views are shown. White arrows indicate the loss of *runx1* expression in PLM. Scale bars are 500 μm. The numbers in the lower left hand corner indicate the number of embryos with similar expression patterns. **(G-J)** Confocal images of cell cycle progression in wild type (G,I) and *stag1b^-/-^*; *stag2b^-/-^* (H,J) embryos at 48 hpf stained with anti-α-tubulin (cyan), anti-phH3 (yellow) antibodies and Hoechst (magenta). Images are maximum intensity projections of 3 (0.15 μm) optical sections taken from the tail region of 48 hpf embryos. Scale bars are 5 μm.

The gene encoding haematopoietic and neuronal transcription factor Runx1 is expressed in the anterior lateral plate mesoderm (ALM), the posterior lateral plate mesoderm (PLM), and in Rohon-Beard (RB) neurons in early zebrafish development (Fig. 1D) (Kalev-Zylinska et al., 2002). Rad21 is required for *runx1* expression in the PLM (Horsfield et al., 2007). We previously found that *runx1* expression is normal in individual *stag* mutants (Ketharnathan et al., 2020). However, we observed loss of *runx1* expression in the PLM of the *stag1b^-/-^*; *stag2b^-/-^* embryos and retained *runx1* expression in the ALM and RB neurons (Fig. 1E). This resembles changes in *runx1* expression in the *rad21^-/-^* mutant (Fig. 1F) (Horsfield et al., 2007), and is consistent with a requirement for an intact cohesin complex for *runx1* expression in the PLM. In contrast, *runx1* expression was normal in *stag1a^-/-^*; *stag2b^-/-^* embryos (Fig. S1B).

We next examined the morphology of mitotic cells in *stag1b^-/-^*; *stag2b^-/-^* embryos at 48 hpf (Fig. 1G-J). In contrast to wild type embryos (Fig. 1G, I) condensed chromosomes were disorganized and abnormally distributed in *stag1b^-/-^*; *stag2b^-/-^* embryos, (Fig. 1H). Lagging chromosomes failed to properly separate during anaphase, resulting in some chromosomes remaining in cell centers (Fig. 1J). These findings suggest that cells in *stag1b^-/-^*; *stag2b^-/-^* mutants lack functional cohesin by 48 hpf, leading to a mitotic blockade. Individual *stag* mutants (Ketharnathan et al., 2020) as well as the *stag1a^-/-^; stag2b^-/-^* double mutant, are homozygous viable. However, loss of both, *stag1b* and *stag2b* is embryonic lethal and phenocopies the previously described *rad21^-/-^* mutant (Horsfield et al., 2007). Our results suggest that Stag1b and Stag2b cannot be compensated by Stag1a and Stag2a proteins in zebrafish.

### Cell division proceeds normally in early stage cohesin mutant embryos

Loss of cohesin in *rad21^-/-^* homozygotes or *stag1b^-/-^*; *stag2b^-/-^* double mutants has different effects on *runx1* expression compared with viable mutations in *stag* genes. Therefore, we were curious to know whether cell cycle effects owing to cohesin deficiency could be responsible for gene expression changes, including *runx1.* Mitotic catastrophe occurs in embryos lacking functional cohesin at 48 hpf (Fig. 1H,J) (Horsfield et al., 2007). However, 16-somite *rad21^-/-^* homozygotes have sufficient maternally-deposited cohesin to continue growth for another 24 hours (Horsfield et al., 2007). We chose to compare cell cycle progression in *stag2b^-/-^* homozygotes with *rad21^-/-^* homozygotes and heterozygotes to determine if these mutants alter cell cycle progression during early embryogenesis, the stage when *runx1* expression is disrupted.

Using BrdU incorporation to mark S phases and phosphorylated histone H3 (phH_3_) staining to mark G2/M cells, we found that S phase proceeds normally in cohesin mutants (Fig. 2A-F). Moreover, the presence of cells double-positive for BrdU and phH_3_ indicated that cells progressed from S to M phase in cohesin mutant tailbuds (Fig. 2D-F). Flow cytometry showed no significant differences in the proportions of cells in G1 (2n), S (2-4n), and G2/M (4n) phases between cohesin deficient tailbuds and wild type controls (Figs. 2G, S2) with the exception of *rad21^-/-^* embryos, which had significantly reduced numbers of cells in S phase (Fig. S2F, p=0.0317 Mann-Whitney U test). We conclude that cell division proceeds essentially normally, although *rad21^-/-^* embryos had slightly reduced numbers of cells in S phase. This is consistent with previous findings that even when cohesin complex quantity is substantially reduced, there remains sufficient cohesin to progress through the cell cycle during early embryogenesis (Horsfield et al., 2007; Schuster et al., 2015).

**Fig. 2.**
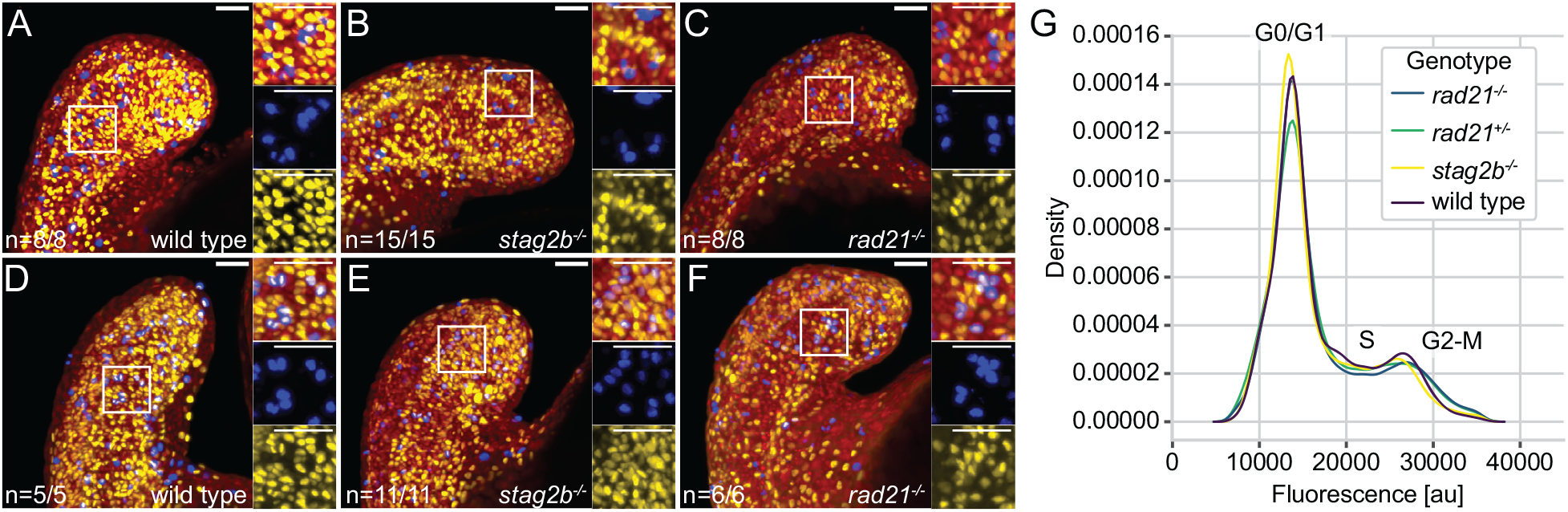
The cell cycle is not blocked in cohesin mutants at the 16-somite stage. **(A-F)** Confocal images showing S phases and M phases in wild type (A, D), *stag2b*^-/-^ (B, E), and *rad21*^-/-^ (C, F) tailbuds at ∼16 hpf. S phases are detected with anti-BrdU (yellow) and M phases with anti-phH3 (blue) antibodies; nuclei are stained with Hoechst (red). BrdU incorporation was measured after incubation for 30 minutes (A-C) or 2 hours (D-F). Zoomed-in images of a selected area indicated by the box are shown. Images are maximum intensity projections of 33 (4.8 μm) optical sections. Scale bars are 40 μm. **(G)** Density plot (*y*-axis) showing the average signal of 3 replicates per genotype over fluorescence signal (DNA stain DRAQ5, *x*-axis; artificial units).

### Cohesin complex quantity and composition affect tailbud gene transcription differently

The zebrafish embryonic tailbud contains neuromesodermal progenitors (NMPs) and midline progenitor cells (MPCs) as well as their neural and mesodermal derivatives (Fig. 3A) and therefore represents an ideal model to study changes in developmental gene transcription and cell fate decisions. Because Stag2 (rather than Stag1) is most likely to be involved in tissue-specific gene transcription (Casa et al., 2020; Cuadrado and Losada, 2020; Kojic et al., 2018; Viny et al., 2019), we compared *stag2b* homozygous mutants with *rad21* homozygotes and heterozygotes to determine how the type of cohesin subunit mutation affects transcription in tailbuds. We performed bulk RNA-seq on 4 pools of 80 excised tailbuds from wild type, *rad21^-/-^*, *rad21^+/-^*, and *stag2b^-/-^* embryos stage-matched at 16 somites. Principal Component Analysis (PCA) separated samples into distinct groups based on their genotype (Fig. 3B). PC1 accounts for 61% of the variance and separated samples into two groups: homozygous and heterozygous *rad21* mutants vs wildtype and *stag2b^-/-^* mutants. PC2 accounts for an additional 11% of the variance and separated *rad21* homozygotes from heterozygotes, and *stag2b* mutants from wild type.

**Fig. 3.**
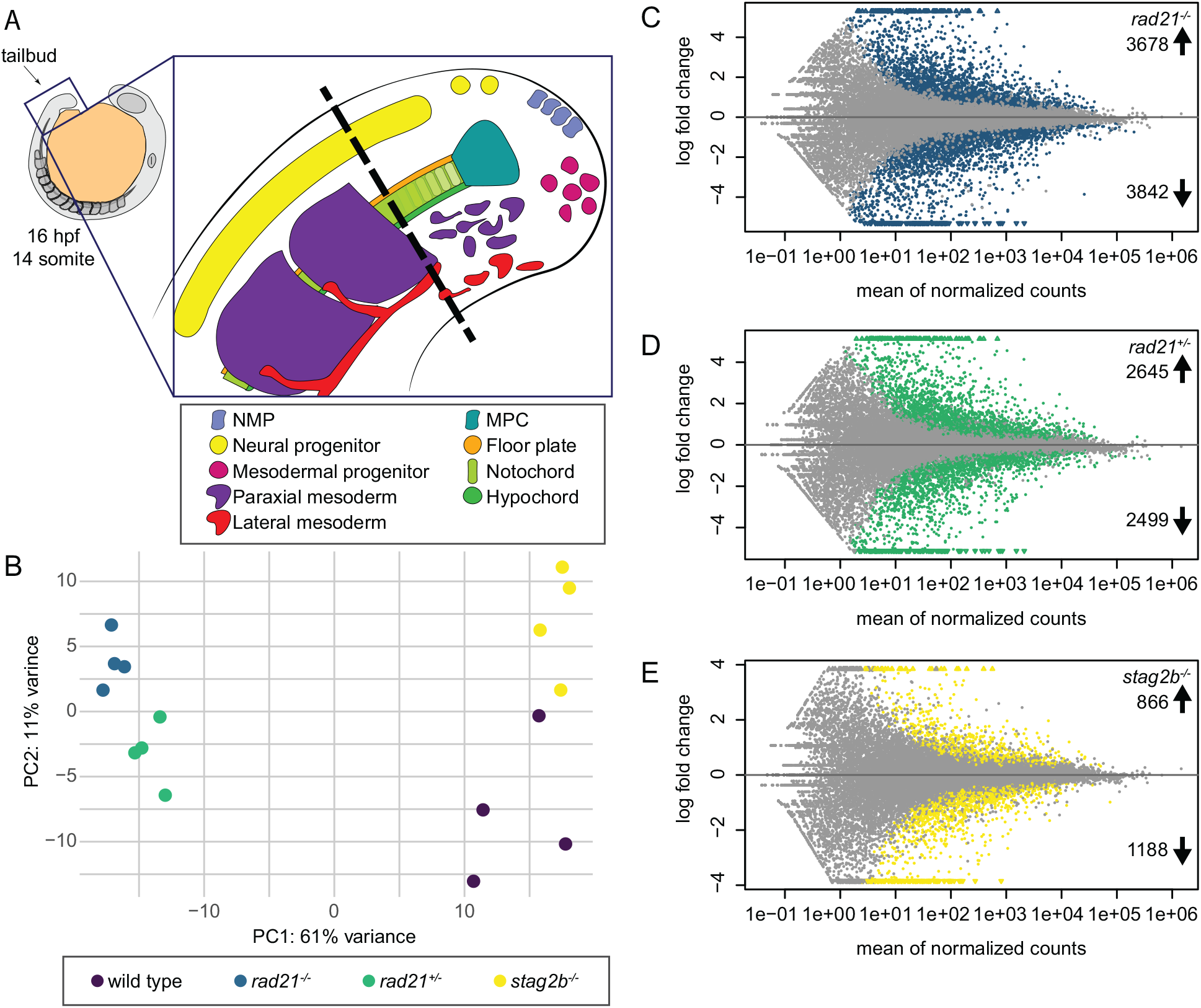
Bulk RNA sequencing analyses of Rad21- and Stag2b-deficient tailbuds. **(A)** Schematic representation of progenitor cells and specialized tissues in the zebrafish tailbud. The zebrafish tailbud consists of two pools of bipotent progenitors: neuromesodermal progenitors (NMPs) and midline progenitor cells (MPCs). The dashed line shows the location of tailbud excision for RNA-seq. **(B)** Principal Component Analysis of gene expression in wild type and cohesin deficient tailbuds at the 16-somite stage. Genotypes are distinguished by colour: wild type samples are displayed in purple, *rad21^-/-^* in blue, *rad21^+/-^* in green, and *stag2b^-/-^* in yellow. **(C-E)** The MA (M (log ratio) and A (mean average) scales) plots display changes in gene expression in *rad21^-/-^* (C), *rad21^+/-^* (D), and *stag2b^-/-^* (E) compared to the wild type tailbuds. Each dot represents a gene, with colored dots indicating those with significant (5% false discovery rate, FDR) changes in expression. 7520 genes were dysregulated in *rad21*^-/-^ tailbuds (3678 up- and 3842 downregulated), while 5144 genes were dysregulated in *rad21^+/-^* tailbuds (2645 up- and 2499 downregulated). In contrast, *stag2b^-/-^* tailbuds had substantially fewer dysregulated genes (2054: 866 up- and 1188 downregulated).

Normalized transcript counts of the cohesin subunits in the different genotypes showed that *rad21* mutation is associated with reduced transcript counts of the other cohesin core subunits, *smc1a* and *smc3*, and increased transcript counts of *stag* subunits. In contrast, transcription of core subunits was unaffected or increased in *stag2b* mutants, and *stag1b* transcript counts increased (Fig. S3). The findings are consistent with the idea that *rad21* mutation reduces cohesin quantity while *stag2b* mutation alters cohesin composition. Differential gene expression analysis (Love et al., 2014) revealed that 7250 genes are dysregulated in *rad21* homozygotes (Fig. 3C), 5144 in *rad21* heterozygotes (Fig. 3D) and 2054 in *stag2b* homozygotes (Fig. 3E). Notably, survivable changes in cohesin dose (*rad21^+/-^*) and composition (*stag2b^-/-^*) strongly affect transcription in the tailbud, indicating that normal levels and subunit makeup of the cohesin complex are important for gene expression.

Of the shared significantly dysregulated genes in cohesin mutant tailbuds, 311 were upregulated and 312 were downregulated in all cohesin deficient tailbuds (Fig. S4A,B), with the highest overlap between *rad21^+/-^* and *rad21^-/-^*. Pathway enrichment analysis using Metascape (Zhou et al., 2019) showed that muscle organ development and energy metabolism were upregulated in all three genotypes, with the highest similarity between *rad21^+/-^* and *rad21^-/-^* (Fig. S4C). Of the downregulated gene pathways, none were conserved across all three genotypes, and more pathways were shared between *stag2b^+/-^* and *rad21^-/-^* than with *rad21^+/-^*. A significant number of terms were unique to *rad21^-/-^* tailbuds including regulation of cell fate specification, suggesting possible dysregulation of tailbud progenitor differentiation (Fig. S4D). Pathway enrichment analysis of significantly downregulated genes in *rad21^-/-^* tailbuds using Reactome (Yu and He, 2016) revealed 26 significantly affected pathways, with top hits associated with mitosis and DNA damage repair (Fig. S5). The most affected pathway was cell cycle control, with 165 genes significantly downregulated in *rad21^-/-^*. Although this pathway did not reach a significance threshold in other mutants, 79 cell cycle genes were significantly downregulated in *rad21^+/-^* and 13 in *stag2b^-/-^* (Data S1-3).

The number of shared dysregulated genes between genotypes suggests that transcriptional changes in *rad21^+/-^* mutants more closely resemble *rad21^-/-^* than *stag2b^-/-^* mutants (Fig. S4A-D). Transcriptional changes reflect genotype rather than viability through to adulthood: *rad21^+/-^* and *stag2b^-/-^* mutants are viable and *rad21^-/-^* mutants are not. Additionally, the results suggest that, consistent with the small effect on cell cycle progression in *rad21* mutants during early embryogenesis (Fig. 2), strong transcriptional changes relate to the expression of cell cycle genes in this genotype.

### Subunit-specific effects of cohesin deficiency on transcription in tailbud cell populations

To assess how cohesin deficiency versus composition affects cell fate decisions in the tailbud we used the bulk RNA-seq data to quantify the expression of genes that mark progenitor cells and their derivatives. We used *rad21^-/-^* as a genotype that represents cohesin deficiency and *stag2b*^-/-^ as a genotype that corresponds to altered cohesin composition.

Tailbud NMPs give rise to mesoderm and neuronal fates, while MPCs give rise to floorplate, notochord and hypochord (Fig. 3A). In *rad21^-/-^* tailbuds, genes marking NMPs were upregulated, neural genes were dysregulated (both up- and downregulated) and genes marking all mesoderm fates were downregulated, including lateral mesoderm that may not derive from NMPs (Fig. 4A). In contrast, NMP and mesoderm marker genes were more subtly affected in *stag2b^-/-^* tailbuds, with non-significant downregulation of *tbxta*, *tbxtb*, *tbx16* and *msgn1* (see Fig. S6 for additional expression data). *stag2b^-/-^* tailbuds had increased expression of genes that mark mature somites and decreased expression of genes marking lateral mesoderm (Fig. 4B).

**Fig. 4.**
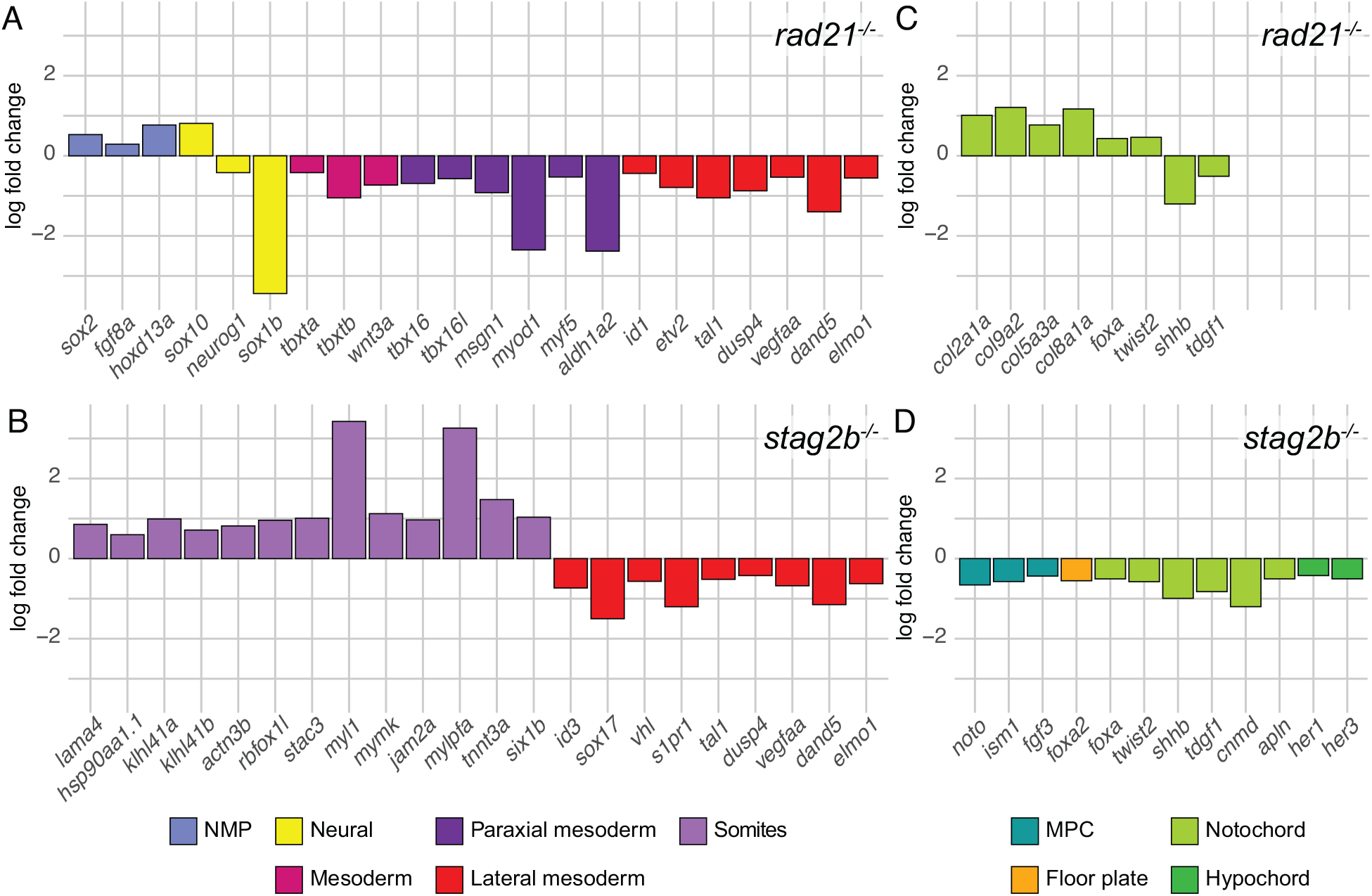
Expression of genes that mark progenitor cells and their derivatives in *rad21* and *stag2b* homozygous mutant tailbuds. **(A-D)** The bar graphs display log2 fold changes significantly (5% FDR) dysregulated marker genes in *rad21^-/-^* (A, C) and *stag2b^-/-^* (B, D) tailbuds compared to wild type. The different categories of marker genes are represented by different colors as specified in the key.

Genes that mark MPCs were expressed normally in *rad21^-/-^*. However, genes encoding notochord-specific collagens were upregulated, and some notochord markers were significantly dysregulated (Fig. 4C). In contrast, genes expressed in MPCs and midline tissues derived from MPCs were significantly downregulated in *stag2b^-/-^* tailbuds (Fig. 4D). The results suggest that *rad21* deficiency causes a block in NMP differentiation, while *stag2b* mutation either affects the composition of mesoderm, or mesoderm gene expression, in tailbuds. Moreover, *rad21* mutation had little effect on midline progenitors (with some effect on MPC derivatives) while *stag2b* mutation reduced transcription of genes expressed in MPCs and all derivatives.

### *rad21* and *stag2b* mutants have different tailbud phenotypes

We next investigated if gene expression changes reflect gross developmental changes in the tailbud in *rad21* and *stag2b* mutants by imaging tailbud cell populations. NMPs are marked by *sox2* and *tbxta* co-expression (Fig. S7). MPCs, a thin band of cells at the end of the notochord, also co-express *sox2* and *tbxta*. Mesoderm progenitors express *tbxta* but not *sox2*, and differentiate into paraxial mesoderm, labelled by *tbx16* expression. *sox2* expression alone labels neural progenitors, lateral mesoderm, the floor plate, and the hypochord (Fig. 3A; Fig. S7) (Steventon and Martin, 2022; Thisse et al., 2001). We used HCR RNA-FISH to visualize the distribution of *sox2*, *tbxta*, and *tbx16* transcripts in cohesin deficient tailbuds (Fig. 5).

**Fig. 5.**
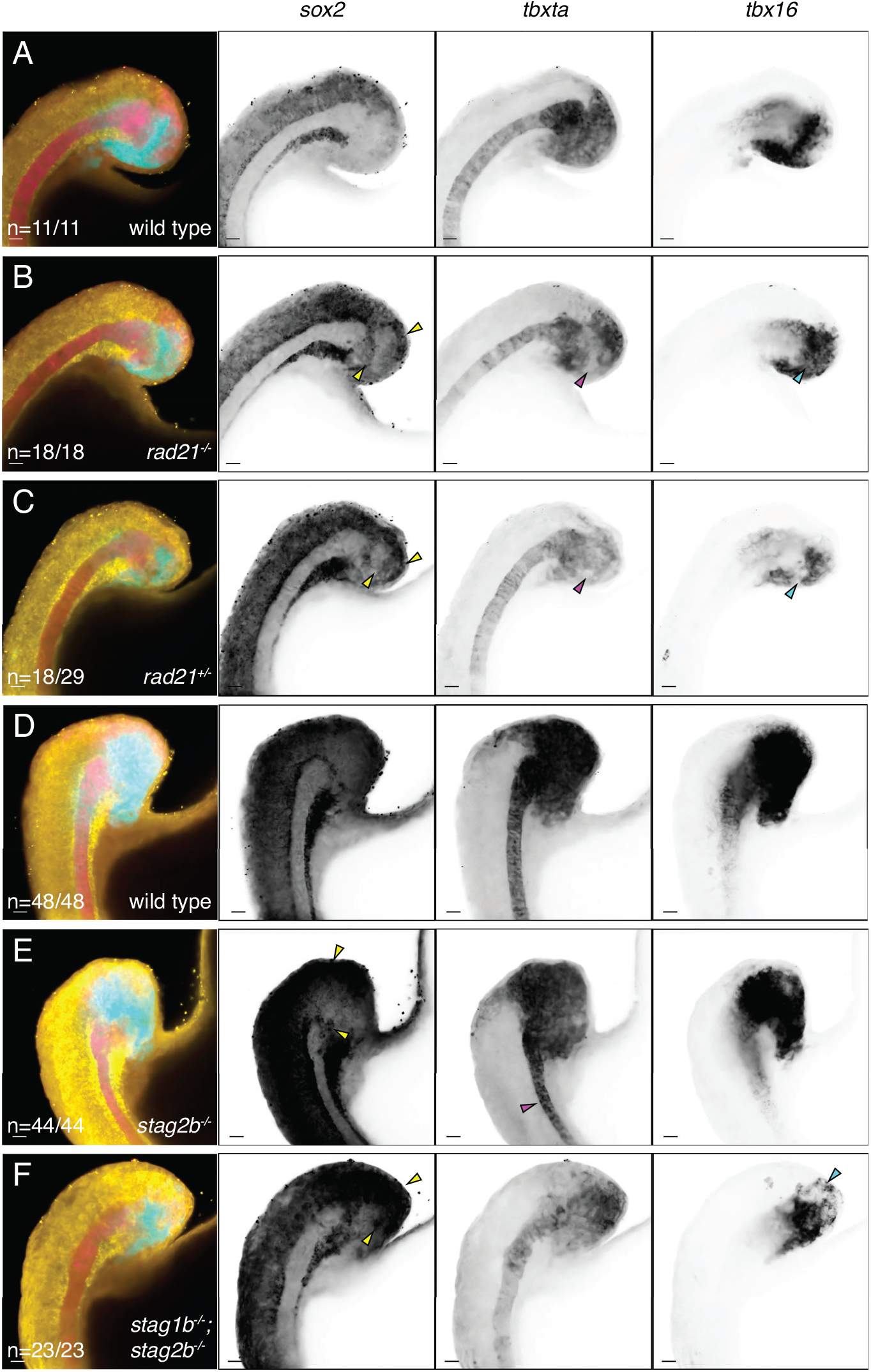
Distribution of *sox2*, *tbxta*, and *tbx16* transcripts in cohesin-deficient tailbuds. **(A, D)** wild type, **(B)** *rad21*^-/-^, **(C)** *rad21*^+/-^, **(E)** *stag2b*^-/-^, and **(F)** *stag1b*^-/-^; *stag2b*^-/-^ zebrafish tailbuds at the 16-somite stage showing expression of *sox2* (yellow), *tbxta* (magenta), and *tbx16* (cyan). Increased *sox2* expression in the NMP region and ectopic expression of *sox2* in the mesoderm is indicated by yellow arrows (B, C, E, F). Pink arrows point to the loss of *tbxta* expression in the region of mesodermal induction (B, C) and the narrow notochord (E), while cyan arrows indicate a decrease in *tbx16* expression. Images are maximum intensity projections of 3 (4.8 μm) optical sections. Scale bars are 20 μm. The number of embryos with each expression pattern out of the total analyzed is noted at the bottom left of the merged panels.

In *rad21^-/-^* homozygotes (Fig. 5B compared with A) and in *stag1b*^-/-^; *stag2b*^-/-^ double mutants (Fig. 5F compared with D) *sox2* expression was expanded at the posterior wall of the tailbud, and the zone of *sox2* expression extended into mesoderm progenitors accompanied by a reduction of *tbxta* expression in these cells. Expression of *tbx16* was restricted to a smaller area than in wild type. Approximately two-thirds of heterozygous *rad21^+/-^* embryos displayed similar expression changes, resembling homozygotes (Fig. 5C compared with A). Like *rad21* mutants, *stag2b^-/-^* mutants had expanded *sox2* expression in the posterior wall of the tailbud (Fig. 5E compared with D). However, in *stag2b^-/-^*, *tbx16* expression appeared normal, while the notochord, visualized by *tbxta* expression, was narrower and did not widen at the posterior end where MPCs reside. Ectopic expression of *sox2* was also observed in this region (Fig. 5E).

We measured the thickness of the notochord (as defined by *tbxta* expression) in wild-type and cohesin-deficient embryos (Fig. 6A,B) and confirmed that notochords were significantly narrower (*p* ≤ 0.0001) in *stag2b^-/-^* embryos (Fig. 6C). In contrast, notochord width in *rad21* homozygous and heterozygous embryos was similar to wild type (Fig. 6C). Surprisingly, *stag1b^-/-^*; *stag2b^-/-^* double mutant embryos had notochords that were normal width (Fig. 5F; Fig. 6C). Therefore, the narrow notochord phenotype was unique to *stag2b^-/-^*, suggesting that the loss of Stag2b impacts MPC differentiation. Because the narrow notochord phenotype is absent in rad21 mutants and stag1b*^-/-^*; *stag2b^-/-^* double mutants, complete cohesin loss is likely epistatic to the narrowed notochord in *stag2b^-/-^* mutants.

**Fig. 6.**
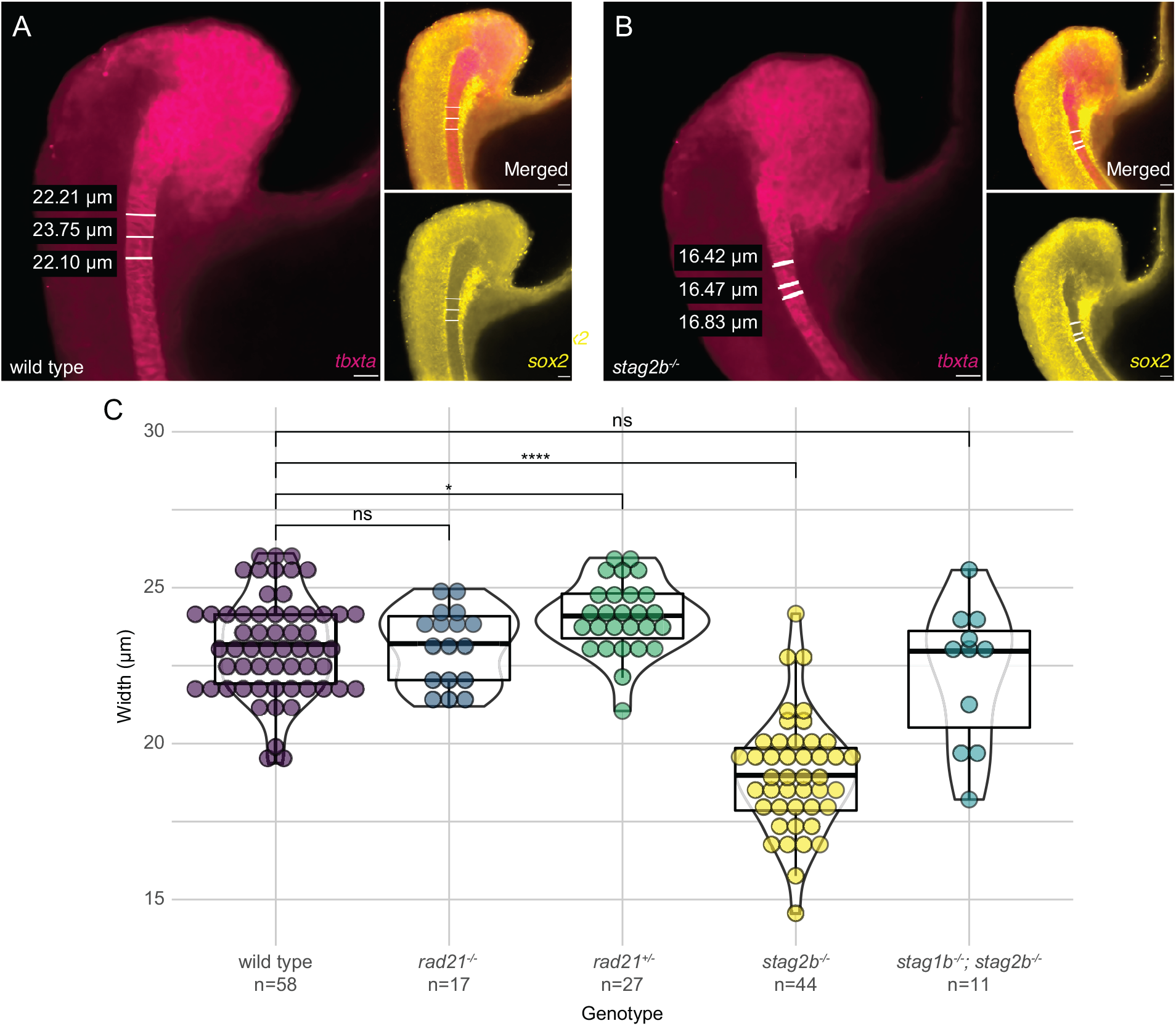
Narrower notochords in *stag2b* mutants are rescued by additional *stag1b* mutation. **(A, B)** Example of notochord width measurement using *tbxta* expression and absence of *sox2* expression. The scale bar is 20 μm. **(C)** Violin plots with overlaid box plots visualizing measurements of notochord width. The genotype and the number of embryos measured in each group are indicated on the x-axis. Significance was determined using an unpaired t-test: * p < 0.05, **** p < 0.0001.

Altogether, our results indicate that different cohesin mutations have different effects on cell populations in the tailbud. Loss of cohesin quantity in *rad21* mutants and *stag1b^-/-^*; *stag2b^-/-^* double mutants caused reduction of *tbx16* and expansion of *sox2* expression, consistent with lack of mesoderm induction. In contrast, *stag2b* mutation (which alters cohesin composition) leads to a narrower notochord.

### Altered cell populations in *stag2b^-/-^* tailbuds likely result from downregulated Wnt signaling in NMPs

It is possible that altered cohesin composition through *stag2b* mutation has unique transcriptional effects on cell fate in the tailbud. We chose to investigate this possibility further using single-cell RNA-sequencing of *stag2b^-/-^* tailbuds compared with wild type at the 16-somite stage.

We integrated the single-cell RNA-seq datasets from wild type and *stag2b^-/-^* tailbuds and annotated clusters representing major cell types based on their gene expression profiles (Fig. S8A). All cell types were present in both wild type and *stag2b^-/-^* samples (Fig. 7A), with minor changes in clusters reflecting altered expression of cell type-specific markers observed in the bulk RNA-seq analysis (Fig. 4C,D). Cell numbers were in slightly different proportions for some clusters (Fig. 7B), with the biggest change being increased numbers of ‘anterior paraxial mesoderm 1’ cells in *stag2b^-/-^* tailbuds compared to wild type (Fig. 7B).

**Fig. 7.**
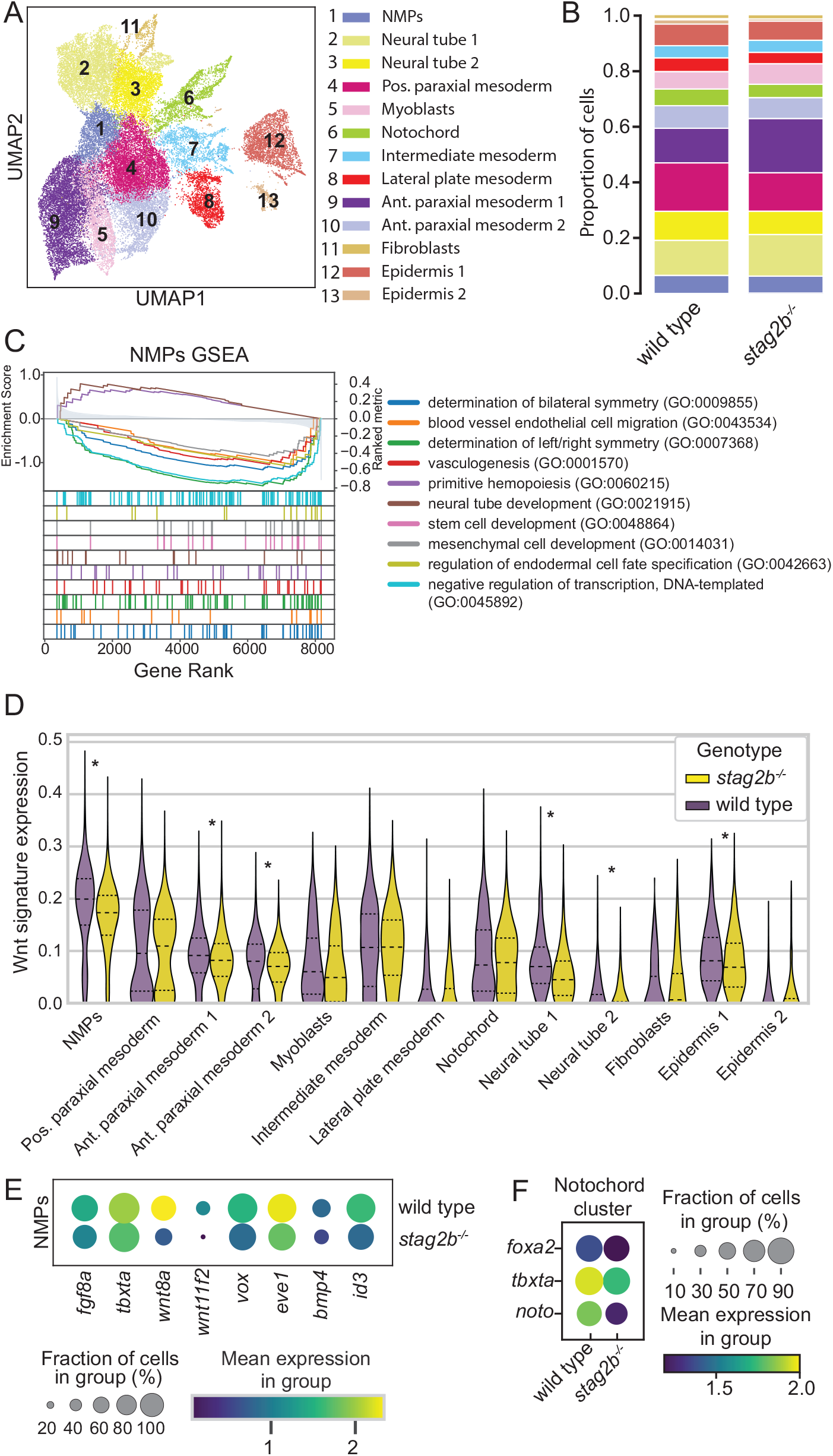
Single cell RNA-seq of tailbuds from embryos at the 16-somite stage shows disruption of Wnt signaling in *stag2b*^-/-^ NMPs. **(A)** Uniform manifold approximation and projection (UMAP) dimensional reduction of two integrated datasets of wild type (15,298 cells) and *stag2b^-/-^* (21,278 cells) tailbud samples (total 36,576 cells) with clustering of the major cell types. **(B)** Stacked bar graph showing cell type proportions inwild type and *stag2b^-/^* **(C)** Gene set enrichments for genes ranked by Z score for differential expression between wild type and *stag2b^-/^* NMP cluster. **(D)** Violin plot of Wnt gene expression signature (log-normalized) among different cell types in *stag2b^-/-^* (yellow) and wild type (purple) embryos. Wilcoxon rank-sum test with 5% FDR. **(E)** Dot plot showing the mean expression of differentially expressed genes part of FGF, Wnt and BMP pathways in the NMP cluster in wild type and *stag2b^-/-^* embryos. (**F**) Dotplot of differentially expressed genes in the notochord cluster.

Curious as to why the anterior paraxial mesoderm would separate into two clusters despite expressing the same lineage genes (Fig. S8A) we further quantified and plotted the expression of cell cycle status genes (Fig. S9A-C). This analysis showed that ‘anterior paraxial mesoderm 1’ is distinct from ‘anterior paraxial mesoderm 2’ by having more cells in G0/G1 and S phase, while all cells are in either G2/M (>90%) or S phase in ‘anterior paraxial mesoderm 2’ (Fig. S9D-F). In *stag2b^-/-^* ‘anterior paraxial mesoderm 1’ cells were more likely to be in G0/G1 phase with fewer cells in S phase compared to wild type. This finding suggests that gene expression or cell population changes in *stag2b* mutants may not be entirely independent of the cell cycle.

NMPs give rise to paraxial mesoderm and to determine if this process is disturbed in *stag2b^-/-^* embryos we performed pseudo-bulk differential gene expression and GSEA (Gene Set Enrichment Analysis) on NMPs and mesoderm clusters (Fig. S10, Data S4). GSEA showed that NMP clusters had downregulated pathways related to stem cell development (GO:0048864), mesenchymal cell development (GO:0048864) and determination of left/right symmetry (GO:0043534) (Fig. 7C, Data S4, S5). These processes are in part positively regulated by canonical Wnt signalling.

We plotted expression levels of Wnt pathway genes across various cell types comparing genotypes (Data S6) and found significant downregulation of Wnt signatures in the NMPs and anterior paraxial mesoderm clusters (Fig. 7D, Data S7). *tbxta* and Wnt ligands (*wnt8a*, *wnt8a-1* and *wnt11f2*) were downregulated in the NMPs, as well as Wnt-responsive genes *vox* and *eve1* (Fig. 7E). In addition there was significant downregulation of *fgf8a* in NMPs suggesting that epithelial-to-mesenchymal transition (EMT) may be deficient in *stag2b^-/-^* embryos (Goto et al., 2017). We also noticed decreased expression of *bmp4* and *id3*; other studies have shown that loss of these genes skews endothelial cell fate towards paraxial mesoderm (Row et al., 2018). Together the results suggest that decreased Wnt signalling and a diversion from endothelial to paraxial mesoderm fate could account for the increased paraxial mesoderm population in *stag2b^-/-^* embryos.

Consistent with the narrower notochord phenotype observed in *stag2b^-/-^* embryos, single cell RNA-seq revealed altered gene expression and cell composition in the notochord cluster (Fig. 7F, Fig. S11A, Data S8). Altered GSEA pathways in *stag2b^-/-^* notochords showed upregulation of Hedgehog signalling, cell adhesion and adherens junctions pathways compared with wild type (Fig. S11B Data S9). Interestingly we observed an increase in hypochord cells in *stag2b^-/-^* tailbuds (Fig. S11C-E, Data S8), which could be occurring at the expense of notochord. Reduced notochord and increased paraxial mesoderm cell numbers in *stag2b^-/-^* could additionally be caused by downregulation of *noto* in *stag2b^-/-^* notochords (Fig. 7F). Consistent with this idea, loss of *noto* causes a switch in cell fates from notochord to paraxial mesoderm in mouse (Yamanaka et al., 2007) and zebrafish (Halpern et al., 1995).

In summary, our results suggest that paraxial and axial midline tissue formation from the NMPs is dysregulated in *stag2b^-/-^* tailbuds. One explanation for these observed changes could be downregulation of Wnt signalling in NMPs that give rise to these tissues.

### Wnt stimulation rescues transcription in *stag2b* but not *rad21* mutant tailbuds

We next determined whether Wnt stimulation could restore transcription in cohesin deficient tailbuds. We performed RNA-seq on tailbuds of embryos treated from shield stage to 16-somite stage with the Wnt agonist, BIO (6-bromoindirubin-3’-oxime), which is a GSK3 inhibitor. Subsequently, we conducted interaction analysis (combined effect of genotype and treatment) (Love et al., 2014) to identify genes exhibiting differential responses to Wnt stimulation in cohesin deficient tailbuds compared to wild type. Heatmaps were used to display clustering of the differentially responsive genes (Fig. 8A-C).

**Fig. 8.**
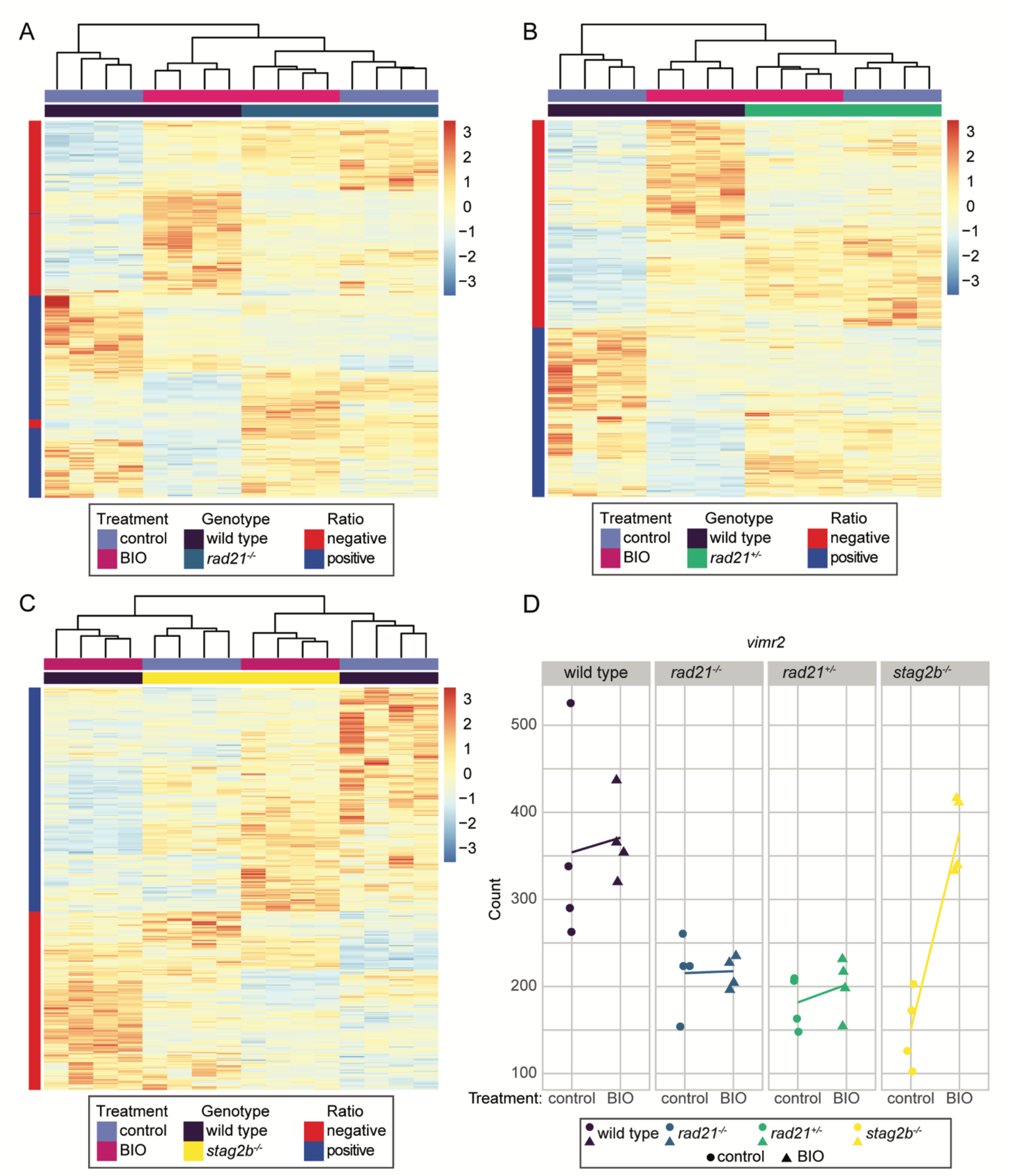
Wnt stimulation normalizes gene expression in *stag2b*^-/-^ but not in *rad21*^-/-^ or *rad21*^+/-^ tailbuds. Embryos were treated from shield stage with 2.5 μM BIO, then tailbuds were collected at 16 somites. 4 replicate pools of 80 tailbuds were used per condition for RNA-seq. The heatmaps display expression levels of the genes that responded differently to BIO stimulation in cohesin mutant genotypes compared with wild type as determined by an interaction analysis. Heatmaps display results from 4 replicates of **(A)** *rad21*^-/-^, **(B**) *rad21*^+/-^, and **(C)** *stag2b*^-/-^ versus wild type. Red and blue colors indicate upregulation and downregulation, respectively, compared to the mean expression. **(D)**, *vimr2* expression is rescued by BIO stimulation in *stag2b*^-/-^ but not in *rad21*^-/-^ or *rad21*^+/-^. Dot plots illustrate the transcript counts of *vimr2* in wild type (purple), *rad21*^-/-^ (blue), *rad21*^+/-^ (green), and *stag2b*^-/-^ (yellow). The x-axis indicates the treatment status, and the y-axis represents the normalised counts. Lines connect the means of the counts for each sample group.

In *rad21^-/-^* and *rad21^+/-^*, the genotype had a stronger effect on clustering of differentially responsive genes than BIO treatment. Genes identified as responding differently to BIO treatment in *rad21* mutants compared to wild type (395 in *rad21^-/-^* and 467 in *rad21^+/-^*) cluster together in the dendrograms regardless of BIO treatment (Fig. 8A,B). Primarily, expression of these genes differs from wild type by being strongly responsive to BIO in wild type, and much less responsive to BIO with homozygous or heterozygous *rad21* mutation. In *stag2b^-/-^* much more complex interactions were observed between the genotype and BIO treatment. In the dendrograms of genes differentially responsive to BIO (539 genes), untreated *stag2b^-/-^* gene sets cluster with BIO-treated wild type, and BIO-treated *stag2b^-/-^* gene sets cluster with untreated wild type (Fig. 8C). This suggests that there is an altered baseline of Wnt signaling in *stag2b^-/-^* and also that BIO stimulation normalizes the expression of select dysregulated genes in *stag2b^-/-^* tailbuds.

A notable example of a gene with expression that is rescued by BIO in *stag2b^-/-^* but not in *rad21* mutants is *vimr2*, a marker of EMT and mesoderm formation in the tailbud (Goto et al., 2017). Expression of *vimr2* was strongly downregulated in all cohesin mutant tailbuds (Fig. 8D, Fig. S12). While BIO treatment had a minimal effect on *vimr2* transcript counts in wild type and *rad21* mutants, it restored *vimr2* levels in *stag2b^-/-^* tailbuds to wild type (Fig. 8D). Interestingly, our single cell RNA-seq data show that *vimr2* is expressed in NMPs and is significantly downregulated in *stag2b^-/-^* mutants (Fig. 9AB). This finding raises the possibility that EMT anomalies marked by downregulated *vimr2* could be responsible for changes in mesoderm induction in *stag2b^-/-^*.

**Fig. 9.**
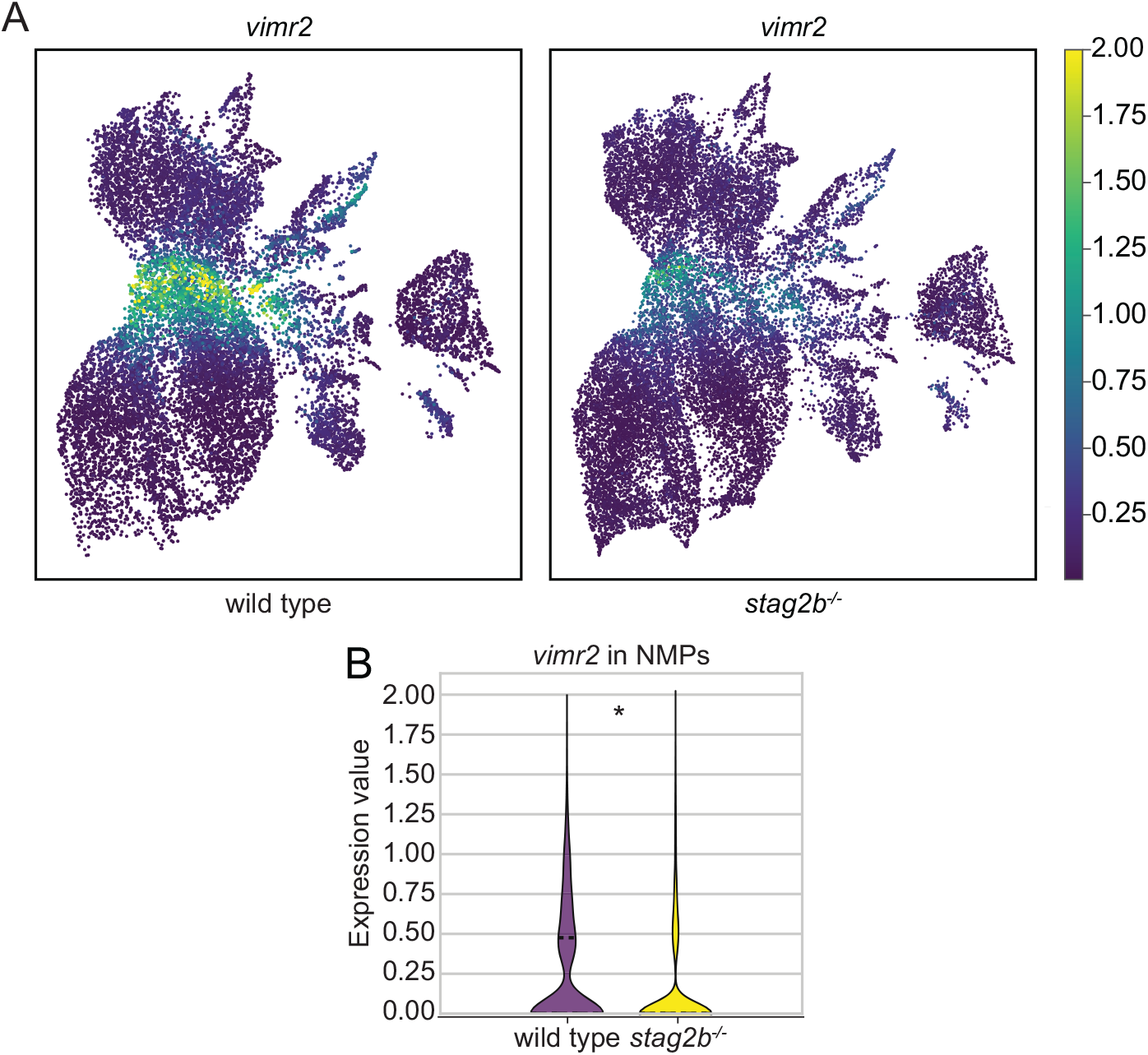
*stag2b* mutation affects *vimr2* expression in NMPs. **(A)** Expression of *vimr2* in UMAP representation in wild type and *stag2b^-/-^* tailbuds at 16 somites. **(B)** Violin plot showing downregulation of *vimr2* expression in the NMPs in *stag2b^-/-^*. * p <0.005, Wilcoxon rank-sum test.

### Wnt stimulation rescues notochord width in *stag2b^-/-^* tailbuds

If BIO stimulation can restore transcription in *stag2b^-/-^* tailbuds, we reasoned that it may also rescue the narrower notochord phenotype in *stag2b^-/-^* embryos. Using HCR with probes for *sox2*, *tbxta*, and *tbx16*, we quantified the thickness of the notochord (*tbxta*) in wild-type and *stag2b^-/-^* embryos both with and without BIO treatment (Fig. 10, Fig. S13). While Wnt stimulation modestly increased notochord width in wild type (*p* ≤ 0.01), it significantly increased the width in *stag2b^-/-^* embryos (*p* ≤ 0.0001) (Fig. 10E). When we compared the notochord width in wild type embryos to that in *stag2b^-/-^* embryos treated with BIO, the difference was statistically insignificant. Therefore, Wnt stimulation rescues the notochord phenotype in *stag2b^-/-^* embryos.

**Fig. 10.**
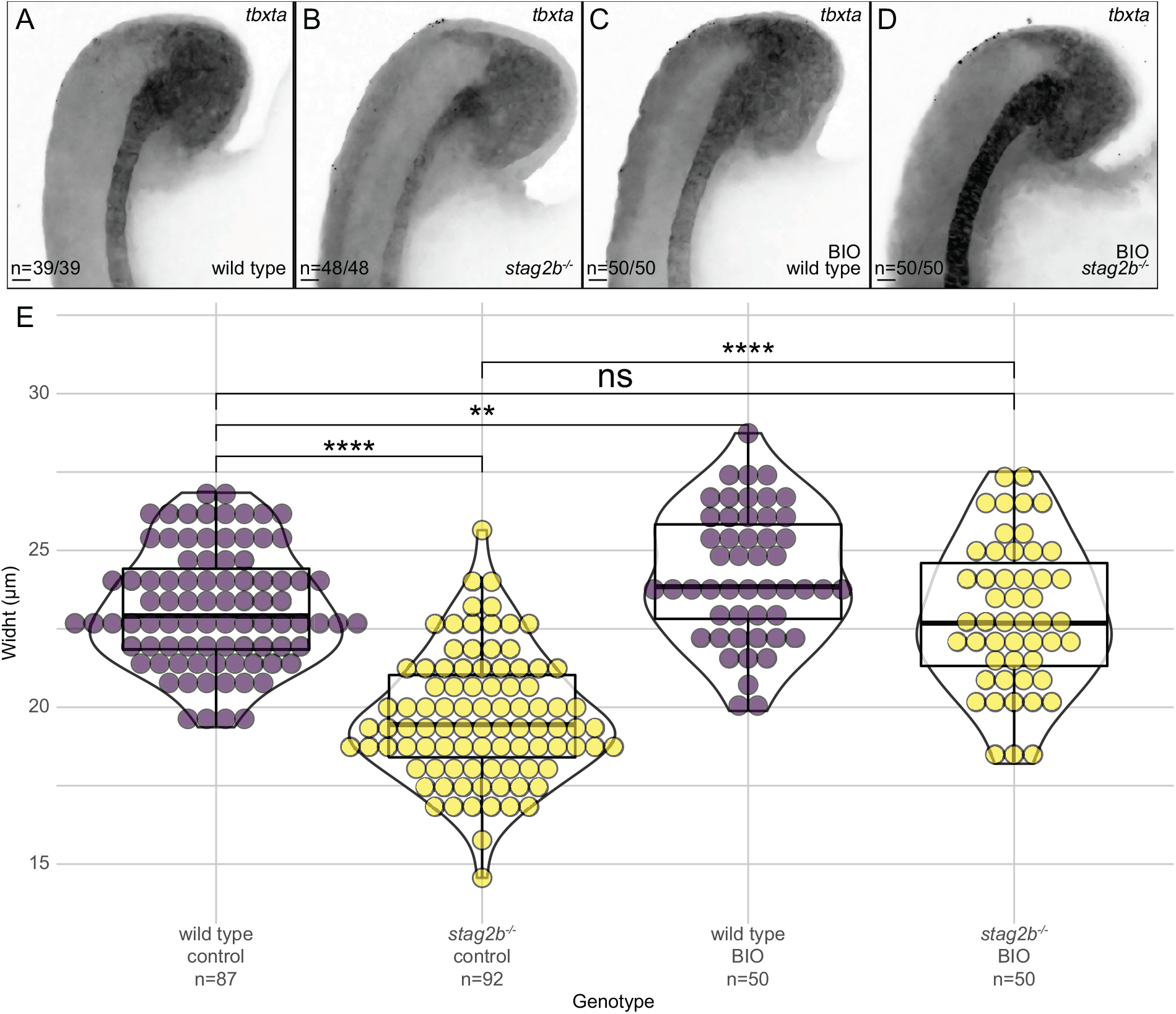
Wnt stimulation rescues notochord width in *stag2b*^-/-^. **(A-D)** Expression pattern of *tbxta* in wild type (A, C) and *stag2b*^-/-^ (B, D) zebrafish tailbuds with (C, D) and without (A, B) Wnt stimulation. Images are maximum intensity projections of 3 (4.8 μm) optical sections. Scale bars are 20 μm. The number of embryos with each expression pattern out of the total analyzed is noted. **(E)** Violin plots with overlaid box plots visualizing measurements of notochord width. The x-axis indicates the genotype, treatment status and the number of embryos measured in each group. Significance was determined using an unpaired t-test: ** p < 0.01, **** p < 0.0001.

In summary, our findings suggest that Wnt stimulation with BIO normalizes gene expression and phenotype in *stag2b^-/-^* embryos, while gene expression in *rad21* mutants is unresponsive to Wnt stimulation.

## Discussion

Germline mutations in subunits of the cohesin complex or its regulators are implicated in developmental disorders known as cohesinopathies, and somatic mutations are now known to cause a variety of cancers. Such cohesin mutations are invariably partial loss-of-function rather than null alleles, because of the essential cell cycle role of cohesin. To date, few studies have compared the developmental consequences of reducing the overall amount of cohesin versus altering its composition. Upon deficiency of Stag2, the Stag1 subunit will compensate in the cohesin complex thereby altering cohesin composition. Upon deficiency of Rad21, the overall quantity of cohesin complexes on DNA decreases. In this study, we took advantage of *stag* and *rad21* gene mutations in zebrafish to show that cohesin composition versus quantity lead to strikingly different consequences for gene transcription and cell differentiation.

Rad21 is an essential subunit in the cohesin complex. Using a zebrafish point mutation *rad21*^nz171^ that progressively reduces *rad21* transcript from heterozygotes to homozygotes, we show that Rad21 deficiency dose-dependently correlates with downregulation of core cohesin subunits, a transcriptional dysregulation signature enriched in cell cycle genes, and a block in mesoderm induction in the tailbud. Although *rad21* heterozygotes are viable and fertile, developmental anomalies in the tailbud are more similar between heterozygotes and *rad21* homozygotes (which die by 48 hpf) than homozygous viable *stag2b^-/-^*. Additional mutation of *stag1b* on top of *stag2b* resulted in loss of viability and a phenotype resembling *rad21* mutants.

It is not clear whether the consequences of Rad21 deficiency are related to cell cycle effects. A recent study showed that blocking the cell cycle in zebrafish does not affect the development of cell types (Kukreja et al., 2023), although it does affect the numbers of presomitic mesoderm cells and erythrocytes (Kukreja et al., 2023). Consistent with cell cycle effects, we observed that *rad21* mutation impacts mesoderm differentiation and our previous work has shown that erythropoiesis is downregulated in cohesin mutants (Horsfield et al., 2007; Ketharnathan et al., 2020). However, cell cycle impairment is unlikely to account for all the defects associated with the reduction in cohesin dose, and it does not explain the transcriptional and phenotypic changes observed in a viable, fertile *rad21* heterozygotes. Our previous work has shown that Rad21 deficiency has remarkably specific transcriptional and developmental consequences, for example, cell-type specific loss of *runx1* expression (Horsfield et al., 2007; Marsman et al., 2014; Schuster et al., 2015). Consistent with a non-cell cycle related transcriptional role for Rad21, complete removal of Rad21 interferes with transcription in post-mitotic neurons, which is rescued upon restoring Rad21 (Weiss et al., 2021). While homozygous mutation in *stag2b* has no statistically significant effect on the cell cycle determined by cytometry, single cell RNA-seq showed an increase in G0/G1 and reduction in S phases in the paraxial mesoderm cell cluster 1 in *stag2b^-/-^* mutants, indicating cell cycle downregulation. It is difficult to determine whether this reduced cell cycle progression in *stag2b^-/-^* mutants is intrinsic, or caused by altered signalling pathways such as Wnt, or a combination of these.

Although *stag2b^-/-^* mutants are viable and fertile, changes in mesoderm differentiation are apparent at tailbud stages. An increase in paraxial mesoderm of *stag2b^-/-^* mutants was detected by both bulk and single cell RNA-seq. Cells over-represented in paraxial mesoderm of *stag2b^-/-^* mutants have exited the cell cycle and are more differentiated, since they primarily reside in the anterior paraxial mesoderm region where somites start to form. We speculate that these characteristics reflect an inability of *stag2b^-/-^* mutants to maintain mesoderm progenitors, thereby lowering the proportion of immature cells relative to mature cells. Consistent with this interpretation, in *stag2b^-/-^* mutants have fewer of the less mature posterior paraxial mesoderm cells and *stag2b^-/-^* NMPs have downregulated stem cell pathways.

*stag2b^-/-^* mutants have a unique, narrow notochord phenotype. An increase in anterior paraxial mesoderm in *stag2b^-/-^* mutants could in turn affect cell fate decisions in the notochord through increased Notch signalling. The single cell RNA-seq data shows an increase in hypochord cell numbers at the expense of notochord in *stag2b^-/-^* mutants, possibly linked to increased Notch signalling from higher numbers of paraxial mesoderm cells (Latimer and Appel, 2006). Reduced Wnt signalling might also directly lead to a decrease in *noto* expression affecting notochord identity. Cells that lose notochord identity were previously shown to end up in the paraxial mesoderm (Halpern et al., 1995; Yamanaka et al., 2007). Finally, EMT driver *vimr2* is downregulated in NMPs and cell adhesion pathways are increased in the notochord, indicating that the ability of cells to undergo EMT and contribute to various developing tissues might be altered in *stag2b^-/-^* mutants. Strikingly, our rescue experiments suggest that driving Wnt signaling can compensate for transcriptional and phenotypic changes in the *stag2b^-/-^* mutants. The noticeably narrower notochord in *stag2b^-/-^* mutants is rescued by stimulation of Wnt signaling via inhibition of GSK3. Moreover, transcription in *stag2b^-/-^* mutant tailbuds is rescued to wild type upon GSK3 inhibition.

We and others have previously reported dysregulated Wnt signaling upon cohesin mutation (Chin et al., 2020; Grazioli et al., 2021; Mazzola et al., 2019; Medina et al., 2016; Pileggi et al., 2021; Schuster et al., 2015) but the directionality of Wnt signaling disturbance remains unclear. We have shown stabilization of *β*-catenin and both up- and downregulation of components of the Wnt signaling pathway, indicating that the effects of cohesin deficiency on Wnt are likely to be complex (Chin et al., 2020). Interestingly, GSK3*α* inhibition was shown to stabilize cohesin on chromatin, promoting continued loop extrusion (Park et al., 2023). Stabilized loop extrusion is dependent on cohesin as it was eliminated with knockdown of Rad21 (Park et al., 2023). The compound (BIO) we used to stimulate Wnt inhibits GSK3 and does not distinguish between *α* and *β* forms. It is possible that GSK3 inhibition was able to rescue transcription and phenotypes in *stag2b^-/-^* mutant tailbuds but not in *rad21* mutants because a reduction in Rad21 reduces the number of complexes that can be stabilized, whereas Stag1b compensates for loss of Stag2b in those complexes.

Stag1-containing cohesin resides primarily at CTCF sites that demarcate contact domains that are invariant between tissues. Stag2-containing cohesin resides at CTCF and non-CTCF sites, where it is thought to regulate tissue-specific transcription (Casa et al., 2020; Cuadrado and Losada, 2020; Kojic et al., 2018; Viny et al., 2019). Viny et al (2019) showed that Stag1-cohesin cannot fully substitute for Stag2 cohesin in haematopoietic stem cells. It is possible that Stag1-containing cohesin has different properties in loop extrusion than Stag2 cohesin (Alonso-Gil et al., 2023; Cuadrado and Losada, 2020) and likely that developmental gene transcription in *stag2b^-/-^* mutant tailbuds is altered because of compensation by Stag1b. We do not believe the other Stag orthologs are major contributors to development in zebrafish owing to the lethality of *stag1b^-/-^*; *stag2b^-/-^* double mutants.

Stag proteins may have functions that are independent of the cohesin complex. For example a recent study found that upon RAD21 depletion, STAG proteins remain bound to chromatin, interact with CTCF, and cluster in 3D (Porter et al., 2023). STAG proteins interact with RNA and R-loops even in the absence of cohesin. Drosophila’s SA cohesin subunit (equivalent to Stag2) is differentially enriched at enhancers and promoters near origins of replication where it is proposed to recruit cohesin (Pherson et al., 2019). In contrast, RAD21 appears to be key for stable binding of cohesin at CTCF sites. A recent study showed that when RAD21 is cleaved, cohesin is released from DNA including at CTCF sites and loops at these elements are lost (Liu and Dekker, 2022). Interestingly CTCF-independent cohesin-anchored loops within chromatin domains persisted despite RAD21 cleavage. The different molecular and structural behaviour of RAD21 and STAG proteins is consistent with the diverse developmental consequences we observed upon germline mutation in these genes.

It is possible that some of the molecular basis for developmental abnormalities is shared between NIPBL deficiency and STAG2 mutation. A study describing single cell RNA sequencing of early-stage mouse embryos with one deleted copy of *Nipbl* showed that these embryos also experience changes in mesoderm fate and have altered mesoderm cell populations (Chea et al., 2024). Nipbl loss altered the regulation of genes involved in EMT, which parallels our findings in *stag2b* mutant zebrafish embryos. This raises the possibility that abnormal regulation of mesoderm fate could be a conserved feature of the cohesinopathies.

Alternate transcriptional and developmental consequences with Stag2 and Rad21 deficiency have implications for the amelioration of cohesinopathies where Wnt agonists have been explored as potential therapeutic agents for individuals with CdLS (Grazioli et al., 2021), and additionally, for the treatment of cohesin-mutant cancers (Chin et al., 2020). Our results suggest that reducing cohesin dose has very different consequences to altering cohesin composition. This indicates that mutations in core cohesin subunits need to be considered differently to mutations in alternate cohesin subunits or cohesin regulators when developing therapeutics.

## Materials and Methods

### Zebrafish Husbandry

Wild type (WIK) (Rauch et al., 1997), *stag1a^nz204^* (Ketharnathan et al., 2020), *stag1b^nz205^* (Ketharnathan et al., 2020), *stag2b^nz207^* (Ketharnathan et al., 2020) and *rad21^nz171^ (Horsfield et al., 2007)* zebrafish lines were maintained at 28 °C according to established husbandry methods (Westerfield, 1995). Zebrafish were housed in the Otago Zebrafish Facility (Department of Pathology, University of Otago, Dunedin, New Zealand). All animal work was performed in accordance with the Otago Zebrafish Facility Standard Operating Procedures (AUP 21-110) and under Environmental Risk Management Authority approval numbers GMC005627, GMD100922 and GMC001366. For all experiments, embryos were developed at 22 or 28 °C.

### Whole-mount *in situ* hybridisations (WISH) and hybridization chain reaction (HCR) RNA-FISH

WISH for *runx1* was performed using 0.5 ng/μL of riboprobe as previously described (Kalev-Zylinska et al., 2002). Probes for *sox2*, *tbxta*, and *tbx16* and HCR reagents were purchased from Molecular Instruments, Inc (USA). HCR was performed according to the manufacturer’s protocol for zebrafish embryos.

### Flow cytometry

Embryos at the 16-somite stage were fixed in methanol (García-Castro et al., 2021) and tailbuds were dissected (n=30). *rad21* heterozygotes and homozygotes were identified by genotyping the heads of individual embryos using genomic DNA extraction (Meeker et al., 2007) followed by a custom TaqMan assay. Cells were filtered through a 40 μm cell strainer and nuclei were stained with DRAQ5 (#ab108410, Abcam) at 5 μM final concentration on ice for 45 min in the dark. Cell cycle profiles of three independent replicates for each genotype were obtained using a BD FACS Aria III (BD Biosciences). Data analysis and plots were generated using Cytoflow (Teague, 2022).

### BrdU incorporation

For cell cycle analyses dechorionated embryos were incubated in 10 mM BrdU in Ringer’s solution for 30 minutes on ice, rinsed three times with Ringer’s solution and incubated for 30 minutes or 2 hours at 28 °C. Embryos were fixed with 4% PFA overnight at 4 °C, dehydrated in methanol and stored at −20 °C in 100% MeOH.

For staining, embryos were rehydrated in a series of 5-minute washes with PBST/MeOH. Embryos older than 24 hpf were treated with 10 µg/ml proteinase K for 10 minutes, followed by three 5-minute washes in PBST and post-fixation in 4% PFA for 20 minutes at room temperature. Samples were then rinsed three times with sterile distilled water. For BrdU staining, embryos were rinsed twice in 2 N HCl and incubated in 2 N HCl for 1 hour at room temperature to denature DNA and expose the BrdU epitope. Alternatively, for antigen exposure, embryos were treated with acetone for 20 minutes on ice.

### Immunohistochemistry

Samples were rinsed twice with sterile distilled water and washed twice with PBST for 5 minutes. Embryos were incubated in blocking solution (0.2% Roche block, 10% FBS, 1% DMSO in PBST) for 30 minutes, followed by a 2-day incubation with primary antibodies at 4 °C. Primary antibodies used are as follows: anti-phH3 (#3377, Cell Signaling Technology; 1:1000), anti-*α*-tubulin (#T6199, Sigma-Aldrich; 1:500), and anti-BrdU (#B35141, Thermo Fisher Scientific). Antibodies were washed off with three 10-minute washes in PBST and two 10-minute washes in 1% FBS in PBST. Embryos were then incubated with secondary antibodies (1:1000) in 1% FBS in PBST at 4 °C for 2 days in the dark. Secondary antibodies used for immunofluorescence were goat anti-mouse Alexa Fluor 488 (1:1000, #A11001, Thermo Fisher Scientific), chicken anti-rabbit Alexa Fluor 647(1:1000, #A21443, Thermo Fisher Scientific). On the second day, Hoechst 33342 (1 µg/ml) (Thermo Fisher Scientific; 1:1000) was added. Embryos were washed five times for 10 minutes with PBST and imaged.

### Microscopy

Fixed embryos were immersed in 70% glycerol to obtain bright field images. Live embryos were anaesthetized with MS-222 (200 mg/L) and embedded in 3% methylcellulose. Bright-field images were captured using the Leica M205FA epifluorescence microscope equipped with a DFC490 camera and Leica Applications Suite software (Leica Microsystems, Germany).

For confocal microscopy embryos were mounted in 1% low melting agarose (w/v). Confocal images were acquired using a Nikon C2 confocal microscope as Z-stacks of the optical sections. The images were processed using NIS-Elements Denoise.ai Software. Maximum intensity projections were used for the figures.

### BIO treatment

30 μM 6-bromoindirubin-3’-oxime (BIO) solution was diluted to 2.5 μM BIO in E3 medium. Embryos were sorted into 50 embryos per plate and treated with 2.5 μM BIO from 4 hpf until tailbud dissection at the 16-somite stage.

### Tailbud bulk RNA sequencing (RNA-seq) and analyses

Tailbuds were dissected from stage-matched embryos at 16 somites (16-18 hpf) as illustrated in Figure 3A. For RNA-seq, tailbuds were individually lysed in 3 μL of RLT + BME (Qiagen RNeasy) and stored in separate PCR tubes at −80 °C to await genotyping of heads (for *rad21^-/-^* and *rad21^+/-^*). Total RNA was extracted from the pools of 80 tailbuds per sample using the RNeasy Micro kit (74104; Qiagen, Germany). Quality and quantity of RNA were assessed using Qubit 4.0 Fluorometer (Thermo Fisher Scientific, USA), Agilent RNA 6000 Nano Kit on 2100 Bioanalyzer (Agilent Technologies, Netherlands) and NanoPhotometer NP80 Touch (Implen GmbH, Germany).

Libraries were prepared from 250 ng of total RNA using the TruSeq Stranded mRNA Library Prep kit (Illumina, USA) and TruSeq RNA CD Index Plate (Illumina, USA) for sample multiplexing. The concentration of the libraries was quantified using a Qubit 4.0 Fluorometer (Thermo Fisher Scientific, USA), and the mean fragment size was assessed using the DNA High Sensitivity KIT on a 2100 Bioanalyzer (Agilent Technologies, Netherlands). 4 nM pooled libraries was sequenced on NovaSeq S1 flow cell by Livestock Improvement Corporation Ltd. (New Zealand).

RNA-seq reads were trimmed using Cutadapt (Martin, 2011), and aligned to the reference genome (GRCz11.98, genome-build-accession NCBI:GCA_000002035.4) with HISAT2 (Kim et al., 2019) and SAMtools (Danecek et al., 2021). FeatureCounts (Liao et al., 2014) was used to generate fragment count matrices. DESeq2 (Love et al., 2014) was used to perform differential gene expression analysis, and multi-testing correction was done using the Benjamini-Hochberg procedure. The false discovery rate (FDR) threshold was set at 5%. Pathway analysis was performed using Reactome (Yu and He, 2016) and Metascape (Zhou et al., 2019). Genotype-specific BIO treatment effects were tested by adding an interaction term (modelling the interaction between treatment and genotype) at the experimental design stage prior to calling differential genes. R package ggplot2 was used for data visualisation (Wickham et al., 2016).

### Single cell RNA sequencing

Stage matched 16-somite embryos (wild type or *stag2b^-/-^*) were dechorionated using pronase (20 mg/ml in E3) and deyolked in calcium-free Ringer’s solution (116 mM NaCl, 2.6 mM KCl, 5 mM HEPES, pH 7.0). Tailbud tissue was dissected using a small needle and pooled (n=30). Tailbuds were then incubated with collagenase (2 mg/ml in 0.05% trypsin, 1 mM EDTA, pH 8.0, PBS) in 1.3 ml volume at 28 °C for 15 mins with intermittent pipetting to achieve a single cell suspension. The reaction was stopped by adding 200 μL of a stop solution (30% calf serum, 6 mM CaCl_2_, PBS). Cells were centrifuged at 500 x *g* for 5 min and re-suspended in 1 ml of resuspension solution (1% Calf Serum, 0.8 mM CaCl_2_, 50 U/ml Penicillin, 0.05 mg/ml Streptomycin). After centrifuging cells were resuspended in 700 μL of resuspension buffer and filtered through a 40 μm cell strainer and kept on ice. Single cell suspensions were processed at the Genomics High Throughput Facility (UC Irvine, USA) according to the manufacturers protocol for the 10X Chromium single cell platform (10X Genomics, Pleasanton, CA. USA) specifically the Chromium Single Cell 3’ Library and Gel Bead Kit v3 (Cat.No. PN-1000128). Libraries were sequenced on a HiSeq2500 platform (Illumina) yielding 1,024,641,718 reads for wild type and 1,166,072,985 reads for the *stag2b^-/-^* sample.

### Single cell RNA sequencing data analysis

Single-cell RNA-seq FASTQ files were demultiplexed using Cellranger (v7.1.0) (Zheng et al., 2017), mapped to the Danio rerio.GRCz11 (danRer11) transcriptome (v4.3.2) (Lawson et al., 2020) including intronic reads. We obtained an estimated number of 18,704 cells (wild type) and 26,360 cells (*stag2b^-/-^*). In the wild type sample mean reads per cell were 54,782, median UMI (unique molecular identifier) were 15,696 and 3,651 genes detected per cell.

In *stag2b^-/-^* we detected 44,236 mean reads per cell, 13,922 median UMIs and 3,561 median genes per cell. The total number of genes detected was 25,879 for wild type and 26,168 for *stag2b^-/-^*. 98.2% and 98.1% of reads (wild type/ *stag2b^-/-^*) had valid barcodes with Q30 of 94.4% for both samples. 92% and 91.5% of the reads mapped confidently to the zebrafish genome. Downstream anlaysis was performed using scvi-tools (v0.20.3) (Gayoso et al., 2022) and scanpy (v1.9.3) (Wolf et al., 2018). After filtering of empty cells doublet removal was performed using Solo (Bernstein et al., 2020). scanpy was used to filter out cells with less than 200 genes, genes detected in less than 3 cells, cells exceeding gene counts of 98% of the median. We also filtered out cells having more than 15% mitochondrial reads (indicating cellular stress).

Filtering and QC steps resulted in 15,298 wild type cells and 21,278 *stag2b^-/-^*cells. The data were then normalised to 10,000 UMIs per cell using the function (scanpy.pp.normalize). Integration of the two datasets was performed using scvi-tools function scVI model (Lopez et al., 2018). Specifically we set up the model using the following command: scvi.model.SCVI.setup_anndata (anndata, layer = “counts”, categorical_covariate_keys=[“Genotype”], continuous_covariate_keys=[’pct_counts_mt’, ‘total_counts’]). We then trained the model and obtained the latent space with model.get_latent_representation(). The neighbourhood graph as well as the UMAP (Uniform Manifold Approximation and Projection) plot was determined using scanpy functions using the scVI latent space as input. Leiden clustering (resolution 0.4) resulted in 13 clusters. Significant differentially expressed genes (DEGs) between clusters and genotypes were determined using the the model calculated above using scVI as well as the Wilcoxon rank-sum test (Benjamini-Hochberg correction).

A list of differentially expressed genes for clusters of interest between wild type and *stag2b^-/-^* tailbuds can be found in Data S4. GSEApy (Fang et al., 2023) was used for gene set enrichment analysis on DEGs identified by pseudobulk analysis and Data S4. We used the function (scanpy.tl.score_genes) for a list of genes obtained from wiki pathways: Canonical_Wnt_pathway (WP1349). The function (scanpy.pp.scale) was used to scale the data to unit variance and zero mean and the Wilcoxon rank-sum test (Benjamini-Hochberg correction) was used to determine significance for the data shown in Fig. 7. Cell cycle phase was determined by using Scanpy’s (Wolf et al., 2018) (sc.tl.score_genes_cell_cycle) function and https://github.com/scverse/scanpy_usage/blob/master/180209_cell_cycle/data/regev_lab_cell_cycle_genes.txt as gene list. Each cell is assigned a score based on the expression of marker genes for S and G2/M phase, if neither gene set score is above a threshold the cell is classified as G0/G1.

### Data availability

Bulk RNA-seq data are available at GEO under accession number GSE247246. Single cell RNA-seq data generated in this study are available at GEO under accession number GSE248952.

## Acknowledgments

The authors would like to thank Noel Jhinku and Dr Doug Mackie for the expert management of the zebrafish facility and Dr Robert Woolley for the confocal microscopy advice. The authors are grateful to Dr Ben Martin and Dr Christian Mosimann for helpful advice and discussions.

## Author contributions

Conceptualization: AAL, JAH

Methodology: AAL, MM, GG, DT, BM, JA, JAH

Investigation: AAL, MM, GG, DT, SK, JAH

Visualization: AAL, MM, GG, DT, SK, TFS, JAH

Supervision: GG, JA, TFS, JAH

Writing—original draft: AAL, MM, JAH

Writing—review & editing: AAL, MM, GG, DT, SK, BM, TFS, JA, JAH

## Supplementary Information

**Fig. S1.**
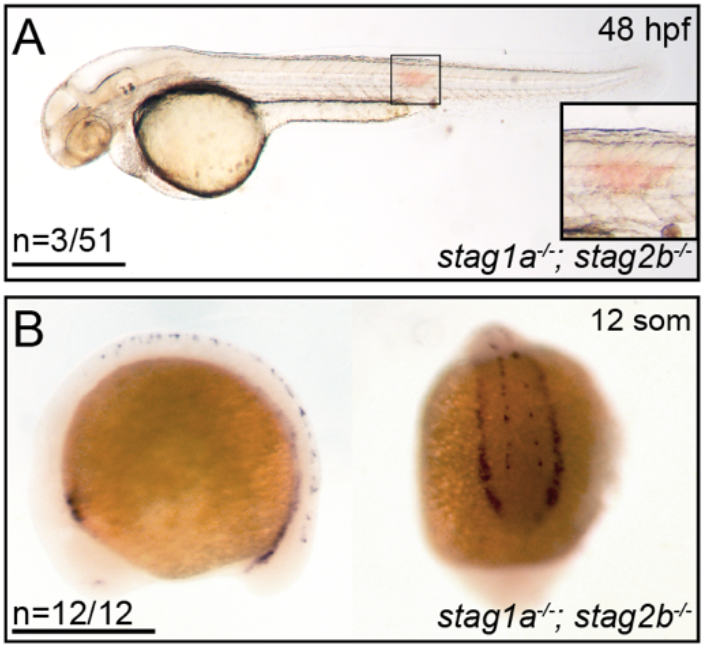
Trunk haemorrhaging and *runx1* expression in *stag1a^-/-^; stag2b^-/-^* mutant embryos. **(A)** Lateral views of representative and *stag1a^-/-^; stag2b^-^*^/-^ embryos at 48 hpf. The boxed region (inset) outlines a trunk haemorrhage representative of those observed in around 5% of the *stag1a^-/-^; stag2b^-/-^* double mutant embryos. Scale bars are 500 μm. **(B)** Normal expression of *runx1* at 12 somites in *stag1a*^-/-^; *stag2b*^-/-^ embryos. Lateral (left) and posterior (right) views are shown. Scale bars are 500 μm. The numbers in the lower left hand corner indicate the number of embryos with the expression pattern shown.

**Fig. S2.**
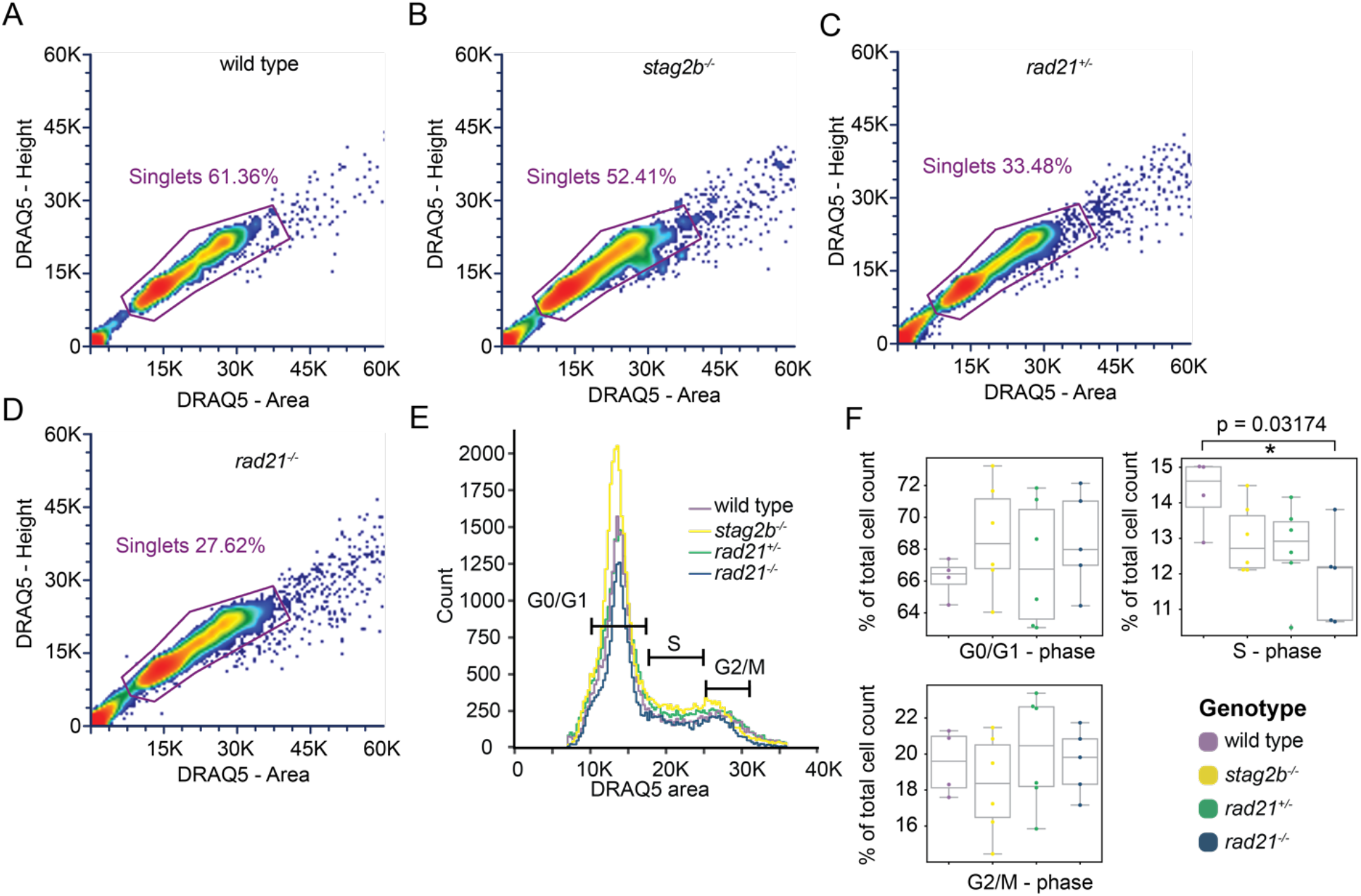
Flow cytometry analyses show that *rad21^-/-^* tailbuds have fewer cells in S phase, while other cohesin-mutant genotypes resemble wild type. **(A-D)** Pseudocolour dot plot of flow cytometry data showing density of cells across DNA content shown as area (*x*-axis) and height of the signal (*y*-axis) of DRAQ5 (labelling DNA) for wild type (A), *stag2b*^-/-^ (B), *rad21*^+/-^ **(C)** and *rad21*^-/-^ (D) tailbud cells. **(E)** Count density plot of wildtype (purple), *stag2b*^-/-^ (yellow), *rad21*^+/-^ (green) and *rad21*^-/-^ (blue) combined replicate samples (n>4). **(F)** Box and swarm plots showing the proportions of cell cycle phases as G0/1, S and G2/M based on DNA content. Boxes show quartiles, whiskers show 1.5 inter-quartile ranges of the lower and upper quartile. *P < 0.05 Mann-Whitney U test.

**Fig. S3.**
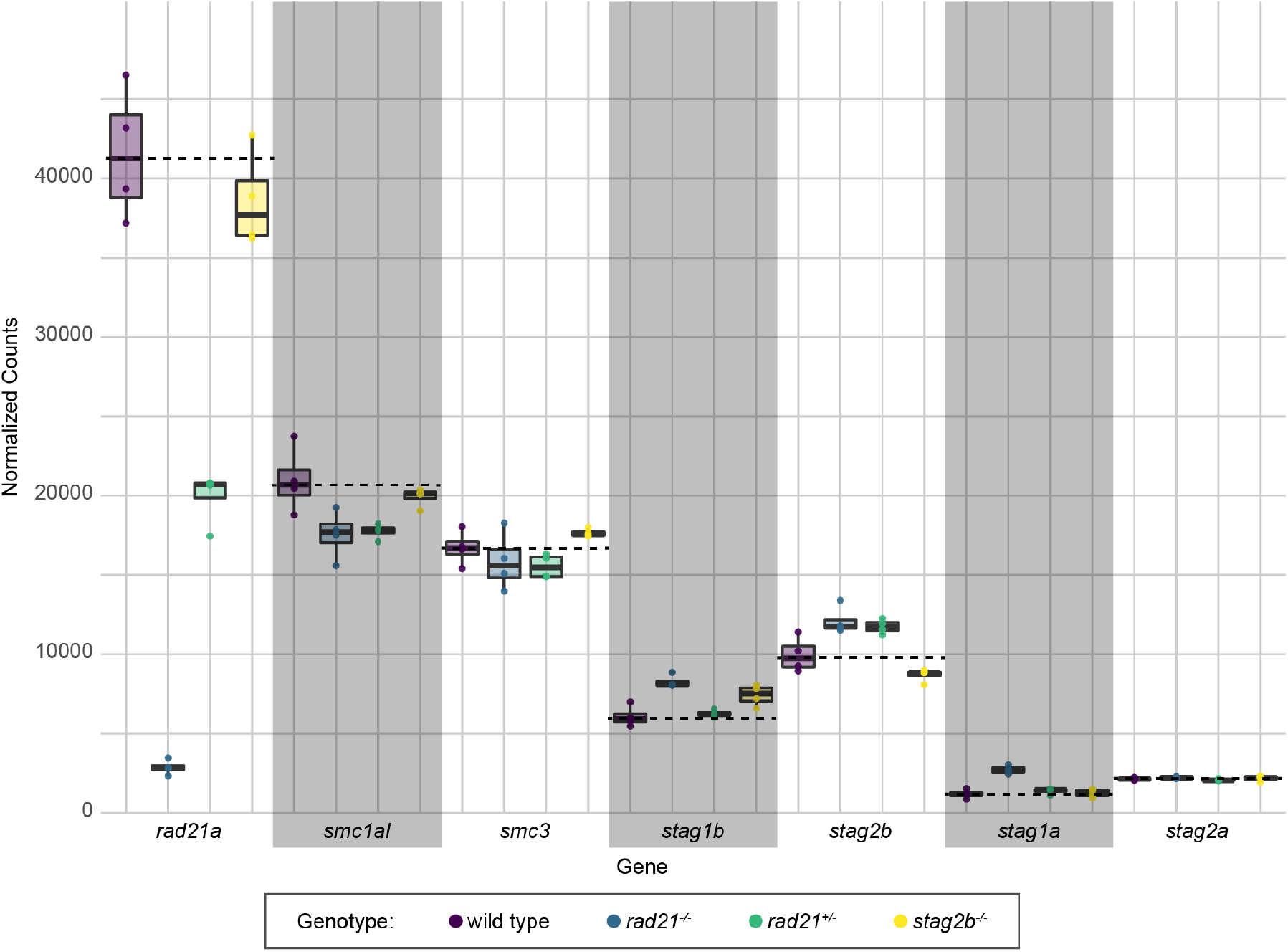
Transcript counts of cohesin subunits in the tailbud. Normalised transcript counts of *rad21a*, *smc1al*, *smc3*, *stag1b*, *stag2b*, *stag1a* and *stag2a* taken from RNA-seq of tailbuds and visualised with box plots defining the distribution of expression levels across these genes in the samples. Genotypes are distinguished by colour: wild type samples are displayed in purple, *rad21^-/-^* in blue, *rad21^+/-^* in green, and *stag2b^-/-^* samples in yellow. For each panel, the dotted line indicates the level of expression in wild type.

**Fig. S4.**
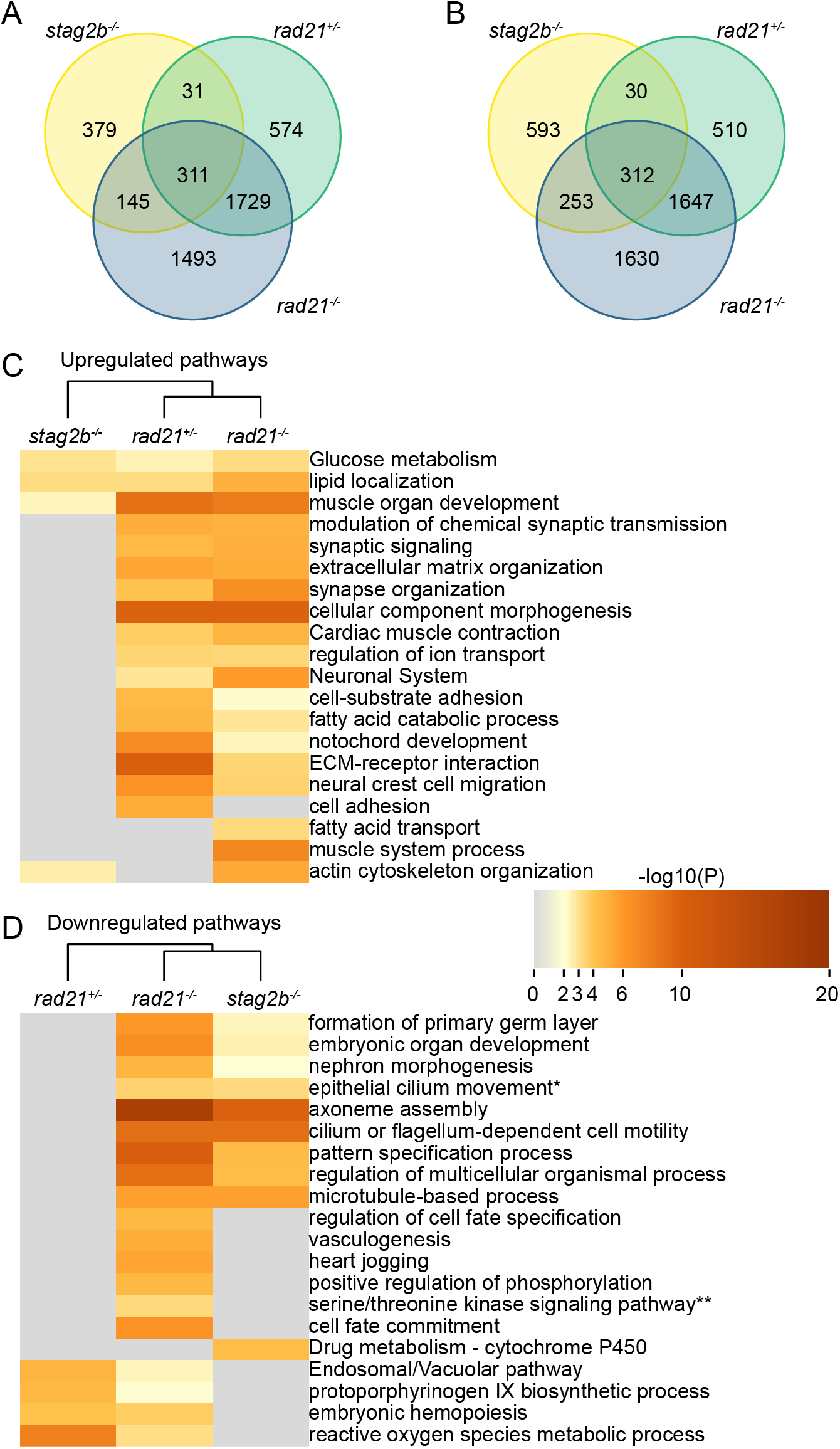
Overlap of dysregulated genes and pathway enrichment in cohesin mutant tailbuds. **(A, B)** The Venn diagrams depict the overlap of significantly upregulated (A) and downregulated (B) genes in cohesin deficient tailbuds. **(C, D)** Metascape heat maps displaying the top 20 terms enriched among significantly upregulated (C) and downregulated (D) genes in cohesin-deficient tailbuds. Corresponding *p*-values are indicated by the colour scale. *Epithelial cilium movement involved in extracellular fluid movement **Regulation of transmembrane receptor protein serine/threonine kinase signalling pathway

**Fig. S5.**
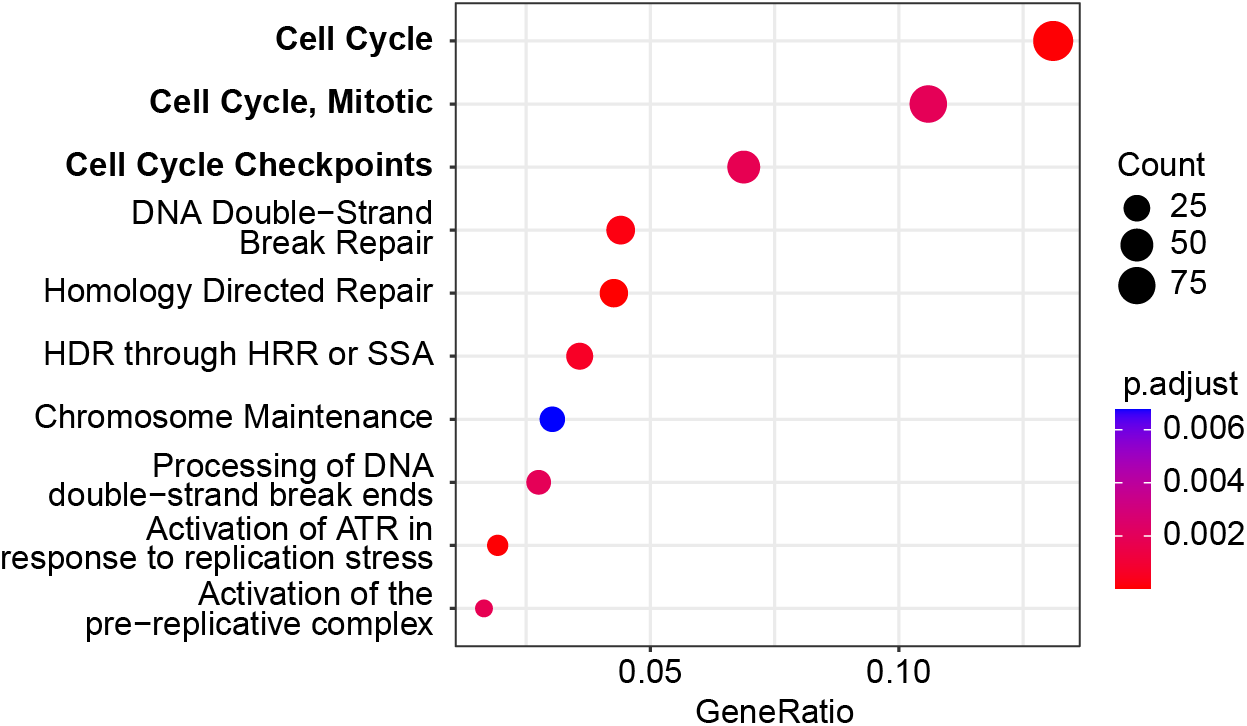
Reactome analyses of downregulated pathways in *rad21^-/-^* mutant tailbuds. The dot plot shows the top 10 enriched Reactome pathways (out of 26) among the significantly downregulated genes in *rad21*^-/-^ tailbuds. The size of each dot indicates the number of genes affected in the pathway, and the dot colour represents the adjusted p-value (Padj). Source data available in Supplementary Data 1-3.

**Fig. S6.**
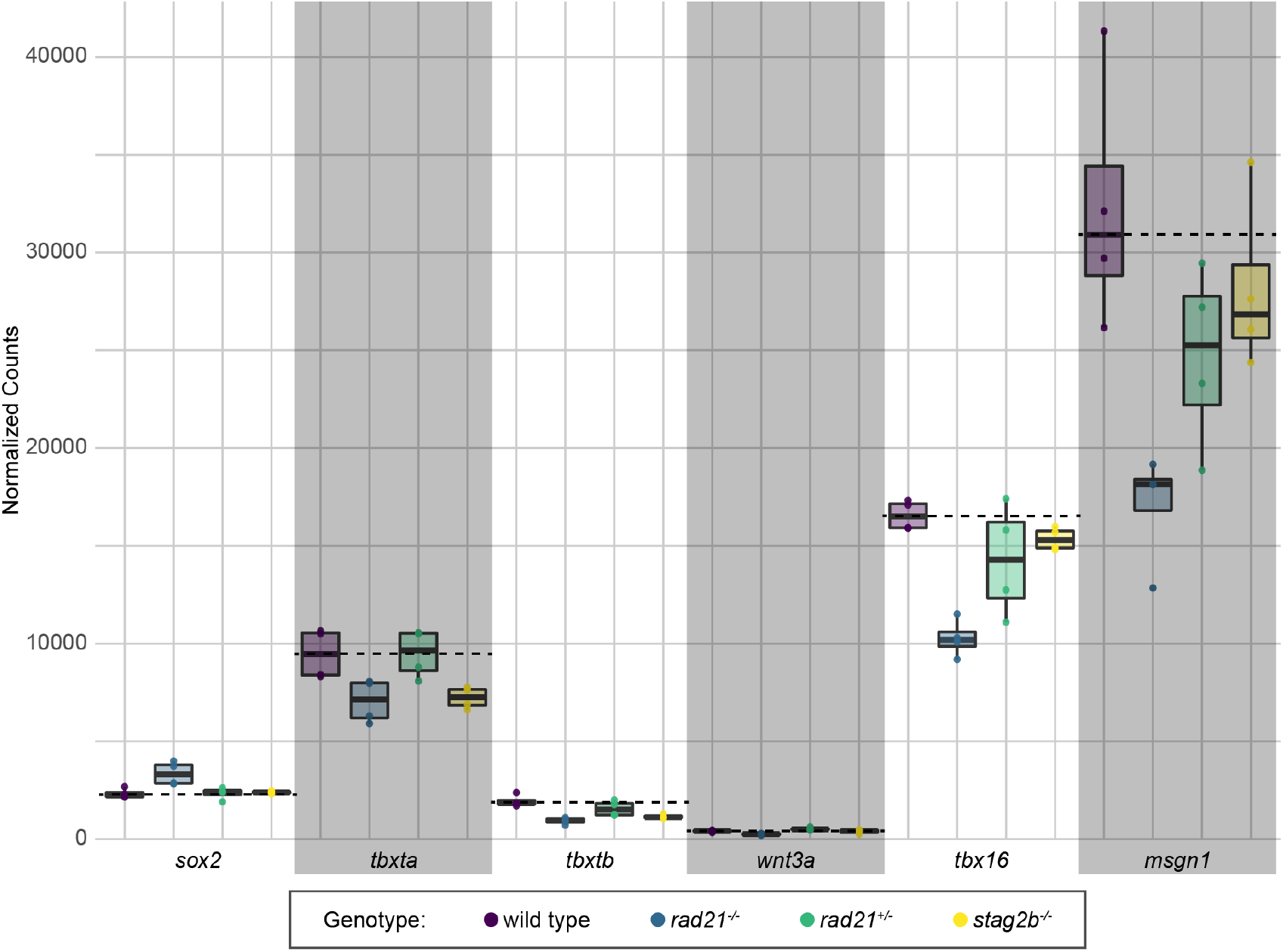
Expression levels of *sox2*, *tbxta*, *tbxtb*, *wnt3a*, *tbx16* and *msgn1* in cohesin mutants. Normalised transcript counts of *sox2*, *tbxta*, *tbxtb*, *wnt3a*, *tbx16* and *msgn1* taken from tailbud RNA-seq data with 4 replicates visualised as box plots defining the distribution of expression levels across these genes in the samples. Genotypes are distinguished by colour: wild type samples are displayed in purple, *rad21*^-/-^ in blue, *rad21*^+/-^ in green, and *stag2b*^-/-^ samples in yellow.

**Fig. S7.**
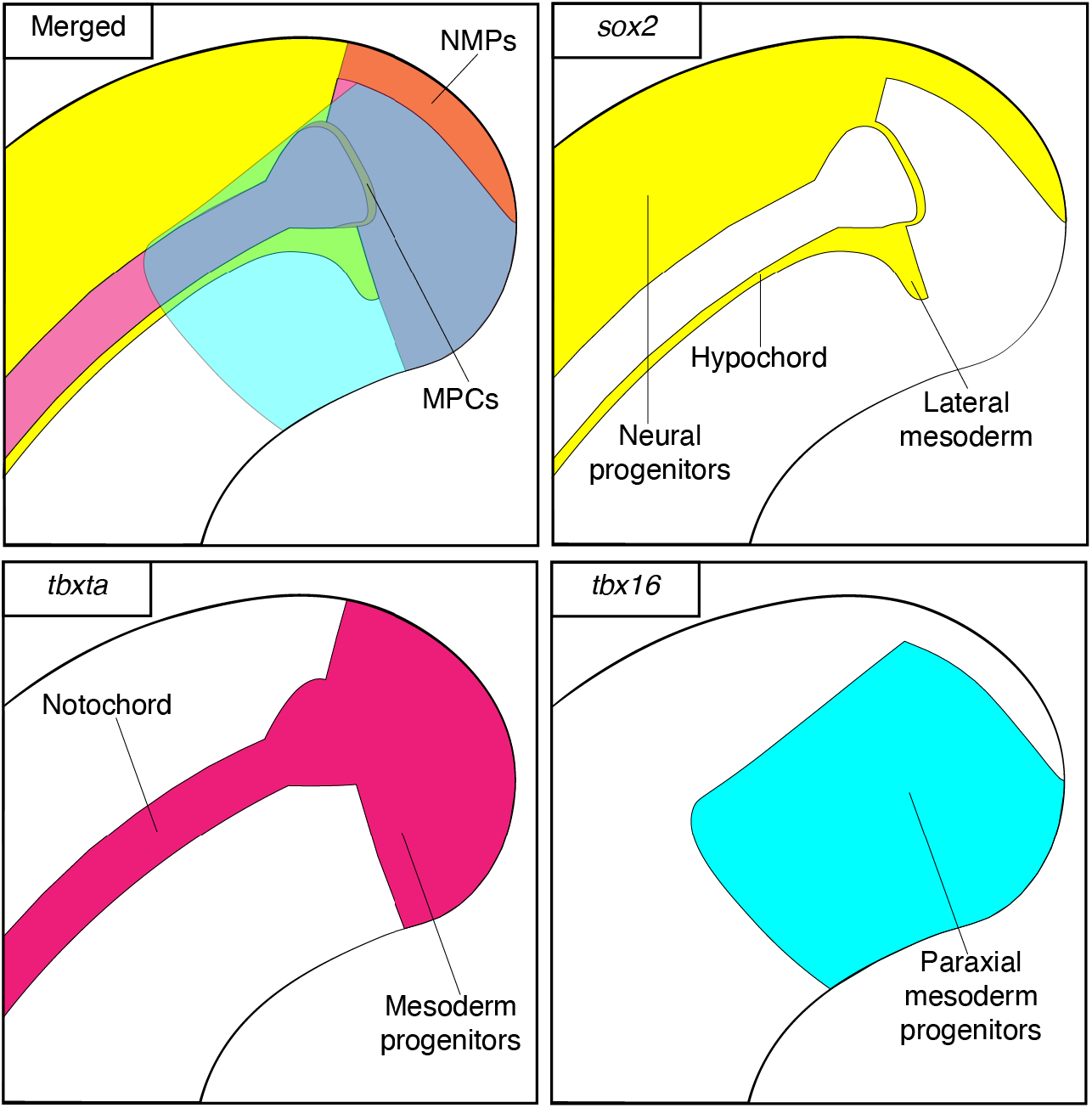
Expression pattern of marker genes in the tailbud at the 16-somite stage. Schematic depicts the regions in the tailbud and progenitor types where expression of *sox2* (yellow), *tbxta* (magenta) and *tbx16* (cyan) expression is expected.

**Fig. S8.**
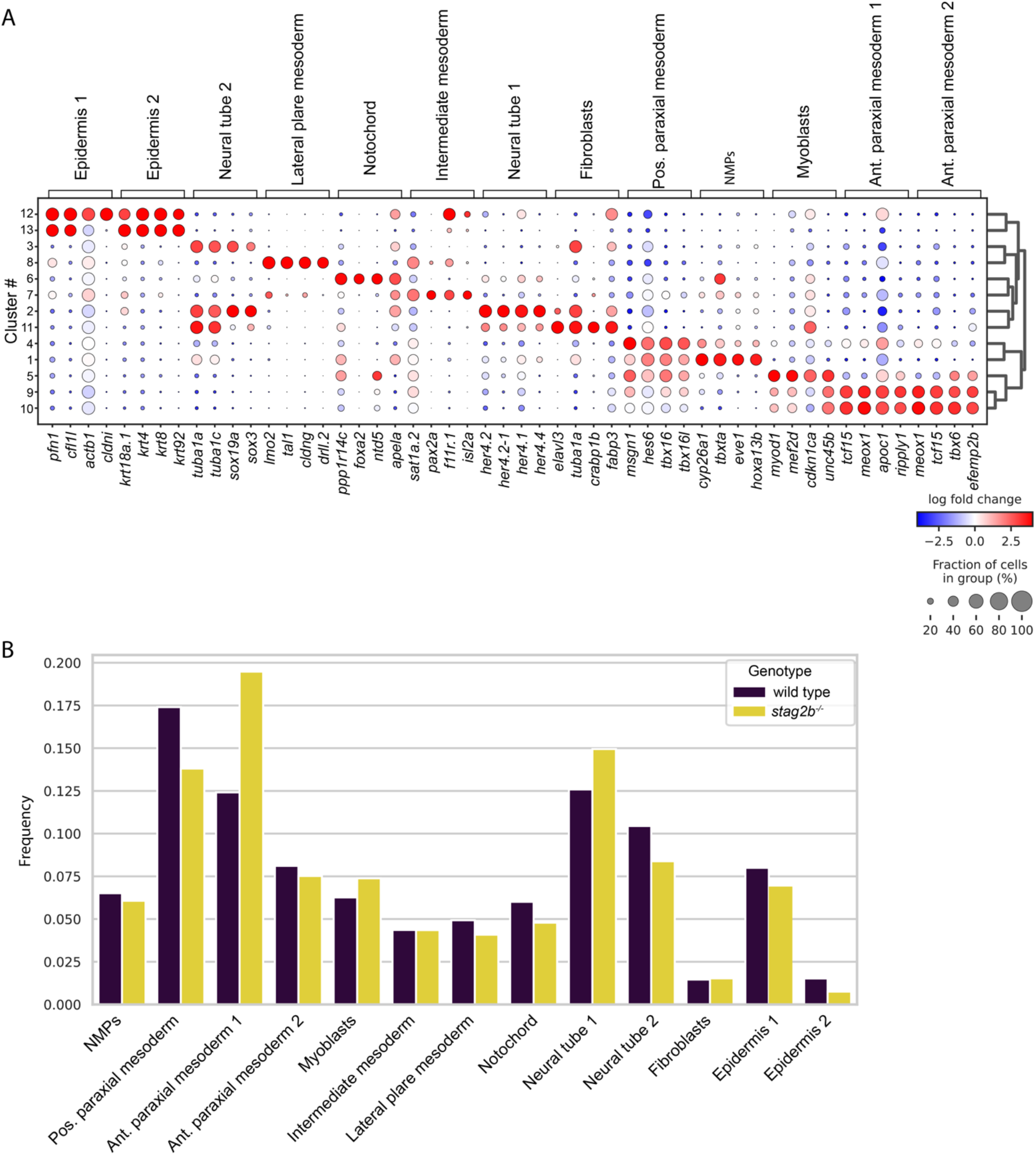
Cell population analysis of single cell RNA sequencing of *stag2b^-/-^* mutant tailbuds compared with wild type. **(A)** Dot plot depicting the expression of the top 4 marker genes per cell population cluster identified (see Fig. 7A). The dot size scales with the fraction of cells expressing the gene, and the dot colour indicates the log fold change between the clusters. **(B)** Differences in the proportion of cell types between wild type and *stag2b*^-/-^ tailbuds among clusters as shown in Fig. 7A.

**Fig. S9.**
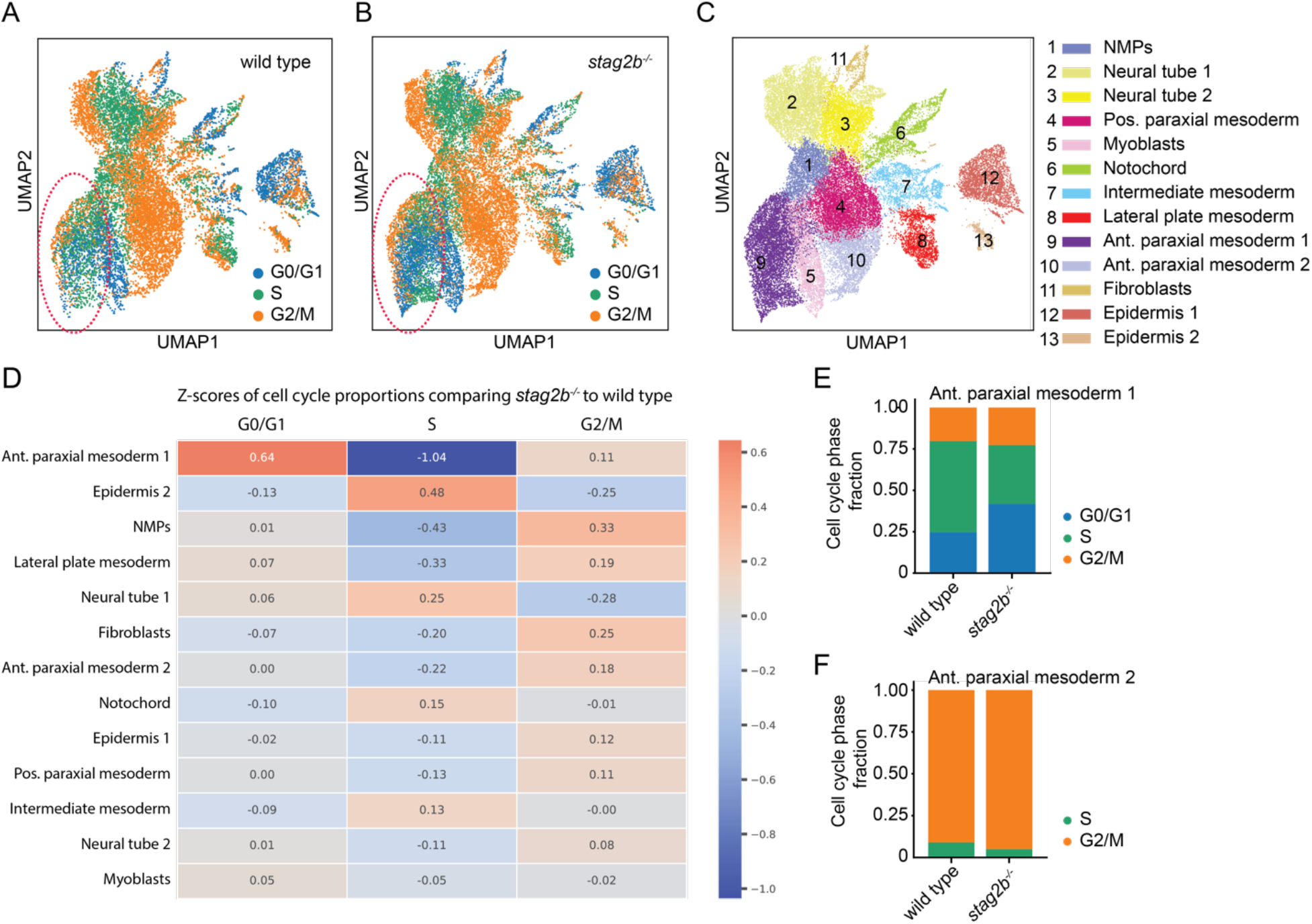
Cell cycle phase analysis of single cell data from *stag2b^-/-^* and wild type tailbuds. **(A-B)** UMAP of wild type and *stag2b*^-/-^ single cell data showing cell cycle phases in blue (G0/G1), green (S) and orange (G2/M). Dashed red outline indicates the anterior paraxial mesoderm cluster 1. All cluster annotations for the integrated dataset in **(C)**. **(D)** Ranked heatmap of z-scores of *stag2b*^-/-^ cell clusters comparing cell cycle phase proportions. Red indicates an increase and blue a decrease in cell cycle phase compared to wild type. **(E)** Stacked bar plot of cell cycle phase fractions comparing anterior paraxial mesoderm 1 cluster in wild type with *stag2b*^-/-^. **(F)** Stacked bar plot of cell cycle phase fractions comparing anterior paraxial mesoderm 2 cluster in wild type with *stag2b*^-/-^.

**Fig. S10.**
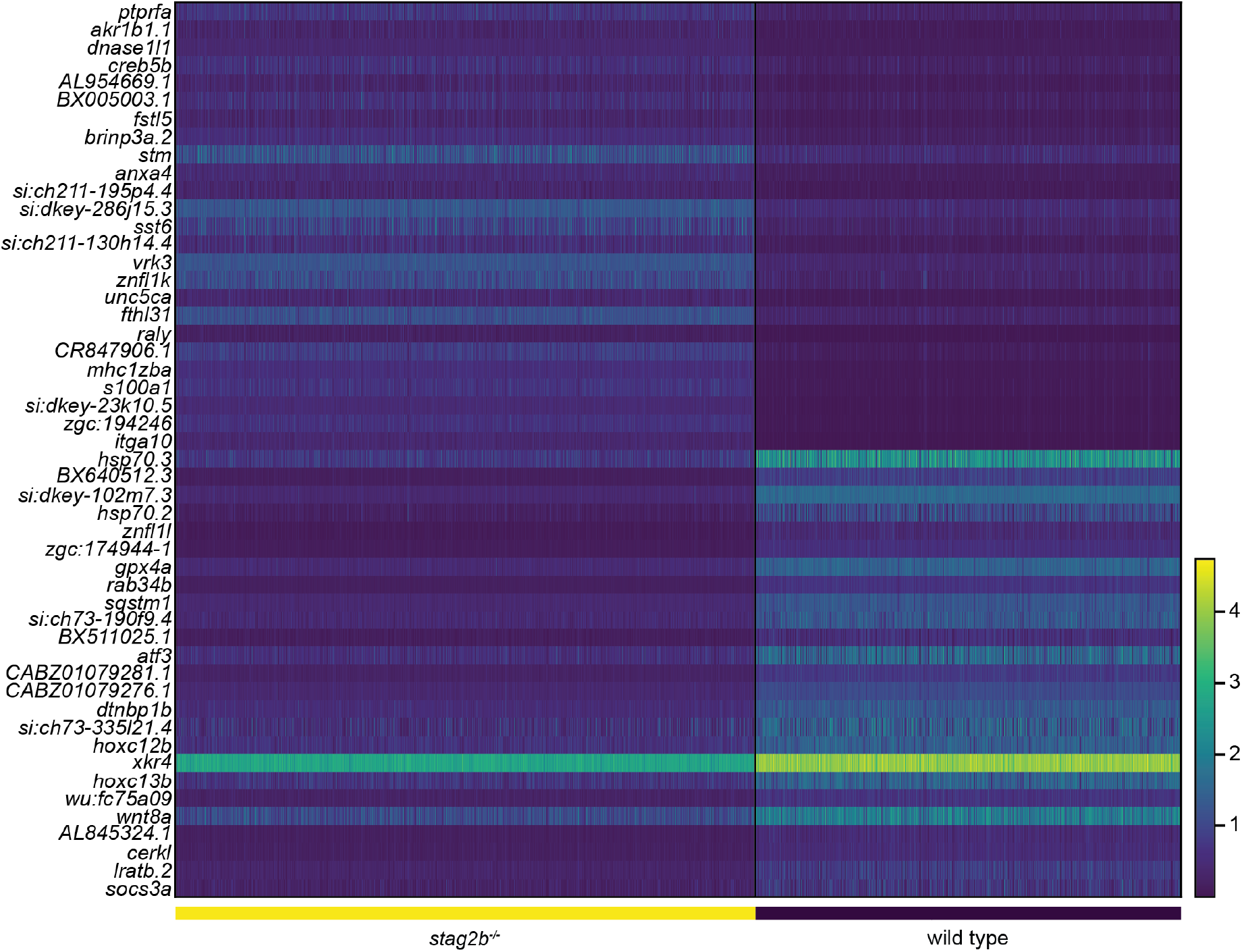
Heat map showing the top 25 differentially up- and downregulated genes between wild type and *stag2b^-/-^* NMP single cell RNA-seq data. For a full list see Supplementary Data 4.

**Fig. S11.**
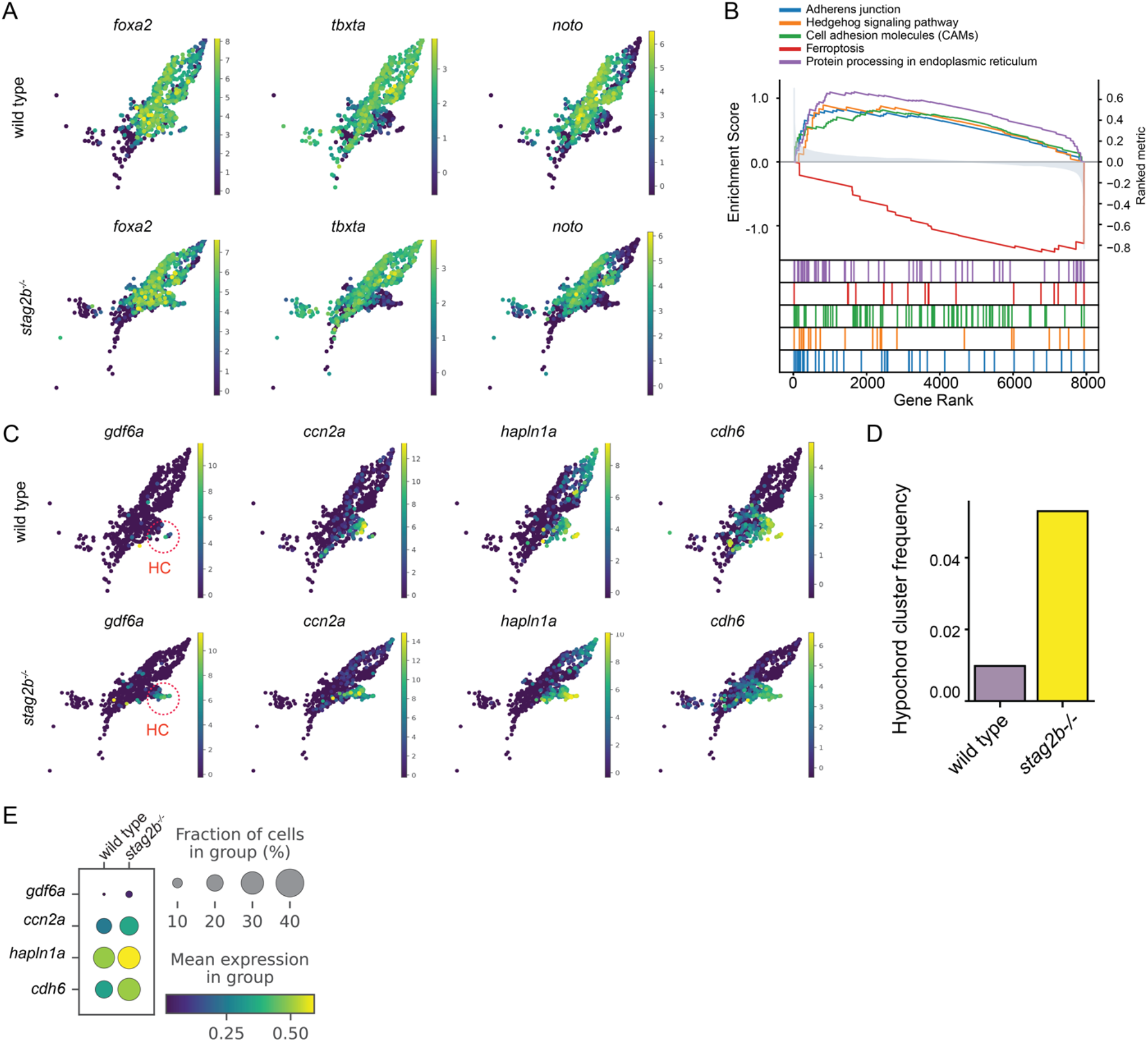
Notochord single cell analysis shows upregulation of Hedgehog signalling with an expansion of hypochord cells in *stag2b^-/-^* tailbuds. **(A)** UMAP plot of notochord markers *fox2a, tbxta* and *noto* showing their expression in wild type (top) and *stag2b*^-/-^ (bottom) notochord cluster. **(B)**. Gene set enrichment analysis of differentially expressed genes derived from pseudo bulk analysis comparing the notochord cluster in wild type and *stag2b*^-/-^ tailbud single cell data. **(C)** UMAP of hypochord markers *gdf6a, ccn2a, hapln1a* and *cdh6* in wild type (top) and *stag2b*^-/-^ (bottom) notochord cluster. **(D)** Bar plot of hypochord-marked cells as proportion of cells in the notochord cluster in wild type and *stag2b*^-/-^ tailbud single data. **(E)** Dot plot of hypochord markers in the notochord cluster comparing wild type and *stag2b*^-/-^ tailbuds.

**Fig. S12.**
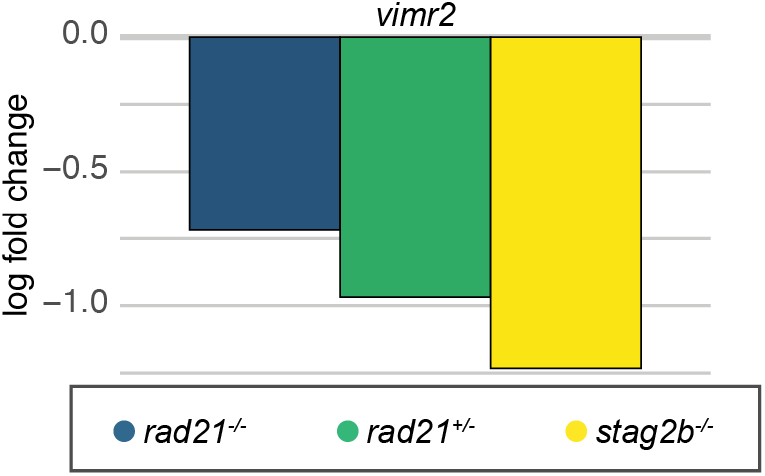
*vimr2* expression in cohesin mutants. The bar graph displays the log2 fold change (5% FDR) for *vimr2* transcripts in *rad21*^-/-^ (blue), *rad21*^+/-^ (green) and *stag2b*^-/-^ (yellow) tailbuds compared to wild type.

**Fig. S13.**
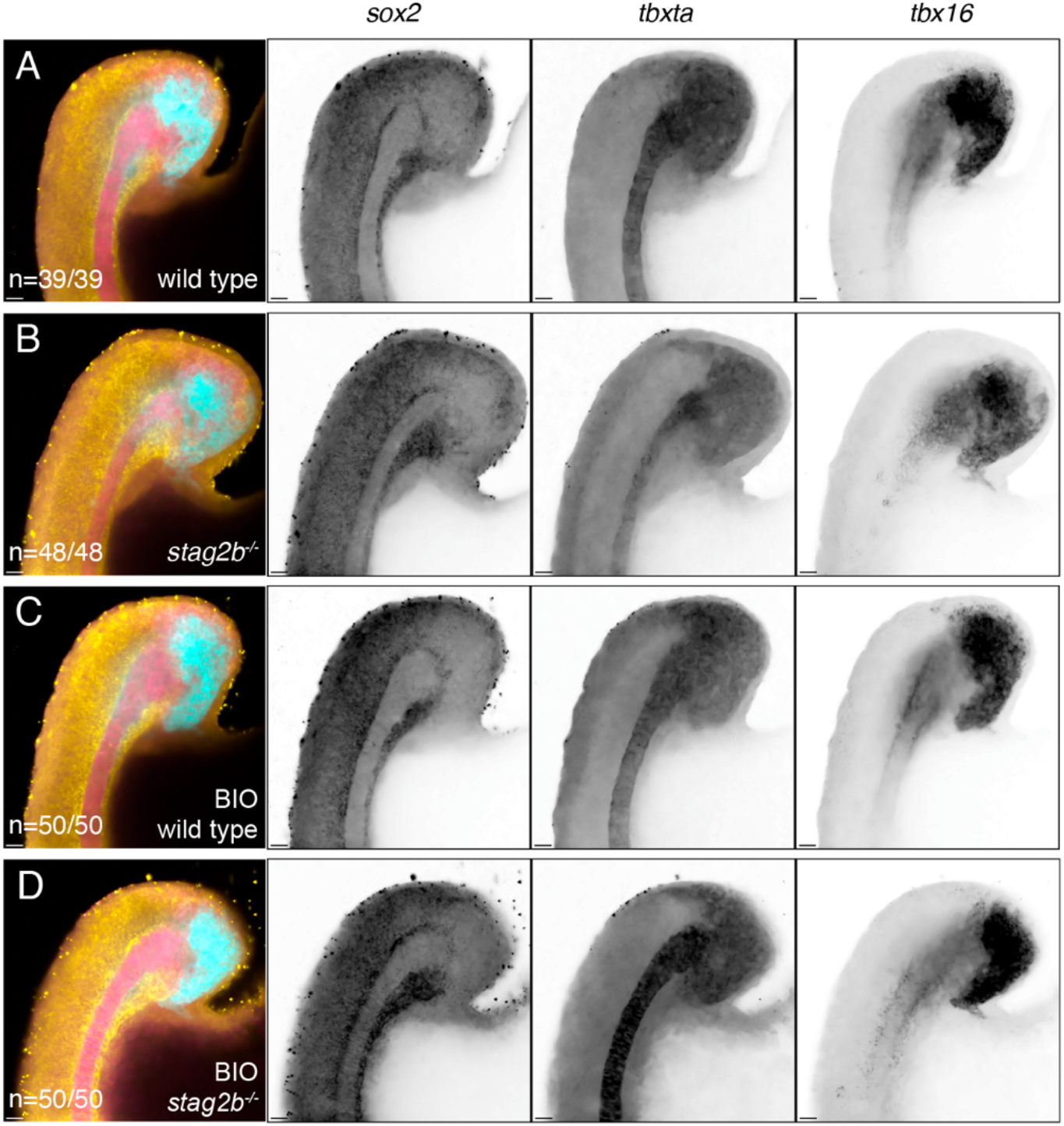
Wnt stimulation rescues the notochord phenotype in *stag2b^-/-^* mutants. **(A-D)** Expression patterns of *sox2*, *tbxta*, and *tbx16* in wild type (A, C) and *stag2b*^-/-^ (B, D) zebrafish tailbuds with (C, D) and without (A, B) Wnt stimulation. Images are maximum intensity projections of 3 (4.8 μm) optical sections. Scale bars are 20 μm. The number of embryos with each expression pattern out of the total analysed is noted.

